# Automated Extraction and Meta-Analysis of a Century of Motor-Unit Research with NeuromechaniX

**DOI:** 10.64898/2026.04.08.717204

**Authors:** Alessandro Del Vecchio, Roger M. Enoka

## Abstract

The scientific literature on human motor units and electromyography (EMG) spans over a century (1925-2025), comprising research impossible to synthesize manually. We introduce NeuromechaniX, a domain-specific platform for automated extraction and meta-analysis of this literature. The core component, MUscraper, is a large language model pipeline that extracts approximately 200 structured metadata fields, organized into 17 major sections spanning participant demographics, EMG acquisition parameters, muscle identification, task protocols, decomposition methods, and motor-unit outcomes, from ∼2,000 publications on human limb muscles. This automated extraction transforms heterogeneous narrative reports into a standardized, queryable database at a scale not achievable through manual review. From this dataset, we analyzed motor-unit discharge rate across 208 studies examining seven muscles. Our analyses reveal that discharge rates differ significantly among muscles (p<0.001), with biceps brachii exhibiting the highest rates (15.9 pps), followed by first dorsal interosseous (13.7 pps) and tibialis anterior (13.5 pps), whereas gastrocnemius (11.3 pps), the vastii muscles (11.5 pps) and soleus show the lowest rates (9.9 pps). Sex-stratified analysis shows females exhibit higher discharge rates than males (14.5 vs 11.9 pps; Cohen’s d=0.38, p=0.018). In contrast, age-stratified analysis reveals non-significant differences between young and older adults (d=-0.24, p=0.072). Collectively, these results show that current views of human motor units are limited to a few muscles, with little data on females and older adults. The complete structured database is available through an open-access interactive platform (https://neuro-mechanix.com/metadata), enabling researchers to explore, filter, and download the extracted metadata. NeuromechaniX provides infrastructure for large-scale meta-research, identification of literature gaps, and hypothesis generation for the neuromechanics community.

## Introduction

Motor units are the functional neural elements through which descending commands and sensory inputs are transformed into muscle force (Sherrington, 1925). Their discharge characteristics provide a direct window into neuromuscular control in health (Enoka, 2008; Kandel et al., 2021; Heckman & Enoka, 2012) and disease (Kimura, 2001), and has been studied extensively using intramuscular EMG (Adrian & Bronk, 1929) and, more recently, high-density surface EMG (HD-sEMG) (Farina & Holobar, 2016; Holobar & Zazula, 2007; Merletti et al., 2008).

The motor-unit literature has grown exponentially over the last century. From the pioneering intramuscular recordings of Adrian and Bronk through the foundational work establishing motor-unit physiology (Basmajian, 1963; De Luca et al., 1982; Bigland-Ritchie et al., 1983; Gydikov & Kosarov, 1974; Milner-Brown et al. 1973; Monster & Chan, 1977), the field has accumulated data gradually over decades. However, the advent of HD-sEMG decomposition has dramatically accelerated the accumulation of these data: by 2020, ∼50 studies using HD-sEMG decomposition had been published (Del Vecchio et al., 2020), and this number continues to increase.

This scale of information fundamentally exceeds the capabilities of manual synthesis. The most comprehensive recent reviews illustrate this limitation: Lulic-Kuryllo and Inglis (2022) systematically reviewed sex differences in motor-unit activity, manually extracting data from dozens of papers, yet this significant effort could only examine one dimension (sex) with a handful of variables from each study. Similarly, other systematic efforts have been constrained to narrow questions: discharge rate modulation (Inglis & Gabriel, 2021), adaptations with aging (Orssatto et al., 2022), and single-muscle analyses (Duchateau & Enoka, 2022). The core problem is that manual extraction, however diligent, can realistically capture relatively few variables from each publication, making it difficult to perform concurrent analyses on discharge rates, muscles, force levels, sex, age, decomposition methods, and electrode configurations. Questions that require cross-tabulating multiple dimensions, such as “Do differences in discharge rate in small and large muscles depend on target force and sex?”, remain unanswerable with traditional review methods.

Two practical barriers limit traditional approaches. First, the study attributes needed for comprehensive aggregation (participant demographics, muscle(s), task parameters, electrode configurations, decomposition methods, and motor-unit outcomes) are reported in heterogeneous narrative and tabular formats that challenge programmatic extraction. Second, retrieval is hampered by inconsistent terminology and, most importantly, many key parameters are scattered across methods, results, and figures.

To address these barriers, we developed NeuromechaniX, a domain-specific platform that combines three components: (1) Automated structured extraction: using MUscraper we transformed 2,331 unstructured publications into structured metadata, enabling aggregation at a scale not feasible through manual methods; (2) Meta-research insights from structured data: the database enables large-scale retrospective analyses revealing muscle-specific discharge characteristics, force-rate relations, and sex differences; (3) A supplementary citation-anchored text summarizer (MUchatEMG; Lewis et al., 2020) that retrieves and summarizes passages from a curated corpus of ∼1,300 peer-reviewed publications, returning citation-grounded responses traceable to specific source papers. However, MUchatEMG is not designed as an alternative to general-purpose language models. Instead, it constrains responses to the peer-reviewed corpus to ensure traceability. The platform enables structured querying and comparative analysis to discover underexplored connections across decades of work.

The main atlas comprises 2,331 selected publications on human limb muscles from 1925-2025, enabling rapid assessment of the distribution of evidence across muscles, tasks, participants, and technologies. We characterize the scope of NeuromechaniX by presenting the two primary contributions: automated structured extraction with MUscraper and meta-research insights from the emergent database. The supplementary text summarization interface (MUchatEMG) is described briefly in the Results and detailed in the Methods.

## Methods

### Literature Search Strategy and Corpus Assembly

We conducted systematic searches across three academic databases to assess the neuromechanics, motor unit, and EMG literature. The search strategy employed 33 structured queries spanning motor-unit decomposition, neural control of movement, and electromyography methodology across PubMed, Scopus, and Google Scholar. PubMed searches combined controlled vocabulary (MeSH terms) with free-text terms covering motor-unit decomposition, discharge rates, recruitment thresholds, conduction velocity, and decomposition algorithms. Scopus queries targeted title, abstract, and keyword fields using Boolean operators to capture HD-sEMG studies, discharge-rate measurements, and signal-processing methodologies. Google Scholar searches specifically targeted review articles and seminal works to ensure comprehensive coverage of foundational literature. The publication date range spanned 1925 to 2025, capturing the full historical record from classical intramuscular techniques through the contemporary HD-sEMG decomposition era.

The final NeuromechaniX corpus comprises 2,331 peer-reviewed publications assembled from the authors’ institutional library access, open-access repositories, and author-provided copies. All publications were obtained through legitimate academic channels and managed via a curated Zotero library, from which bibliographic metadata was exported in CSL-JSON format. A curated subset of 1,300 papers was selected from the full corpus based on availability of full-text access, publication in English, and presence of original motor-unit or EMG data; this subset provides the retrieval corpus for MUchatEMG. Publications span 100 years across 234 unique journals, covering classical neurophysiology, EMG signal processing theory, decomposition methodologies, and contemporary neuromuscular studies.

### Data Flow from Full Corpus to Specific Analyses

As illustrated in Figure 1A, the NeuromechaniX pipeline processes data through multiple stages, with each downstream analysis using a subset of the full corpus based on data availability and inclusion criteria. Table 1 summarizes the sample sizes for each major analysis presented in this work. The full corpus of 2,331 publications yields 30,503 unique participants from 1,774 studies with valid demographic reporting. However, not all studies report all variables; consequently, each specific analysis draws from a different subset of studies depending on which parameters were reported.

**Figure 1.**
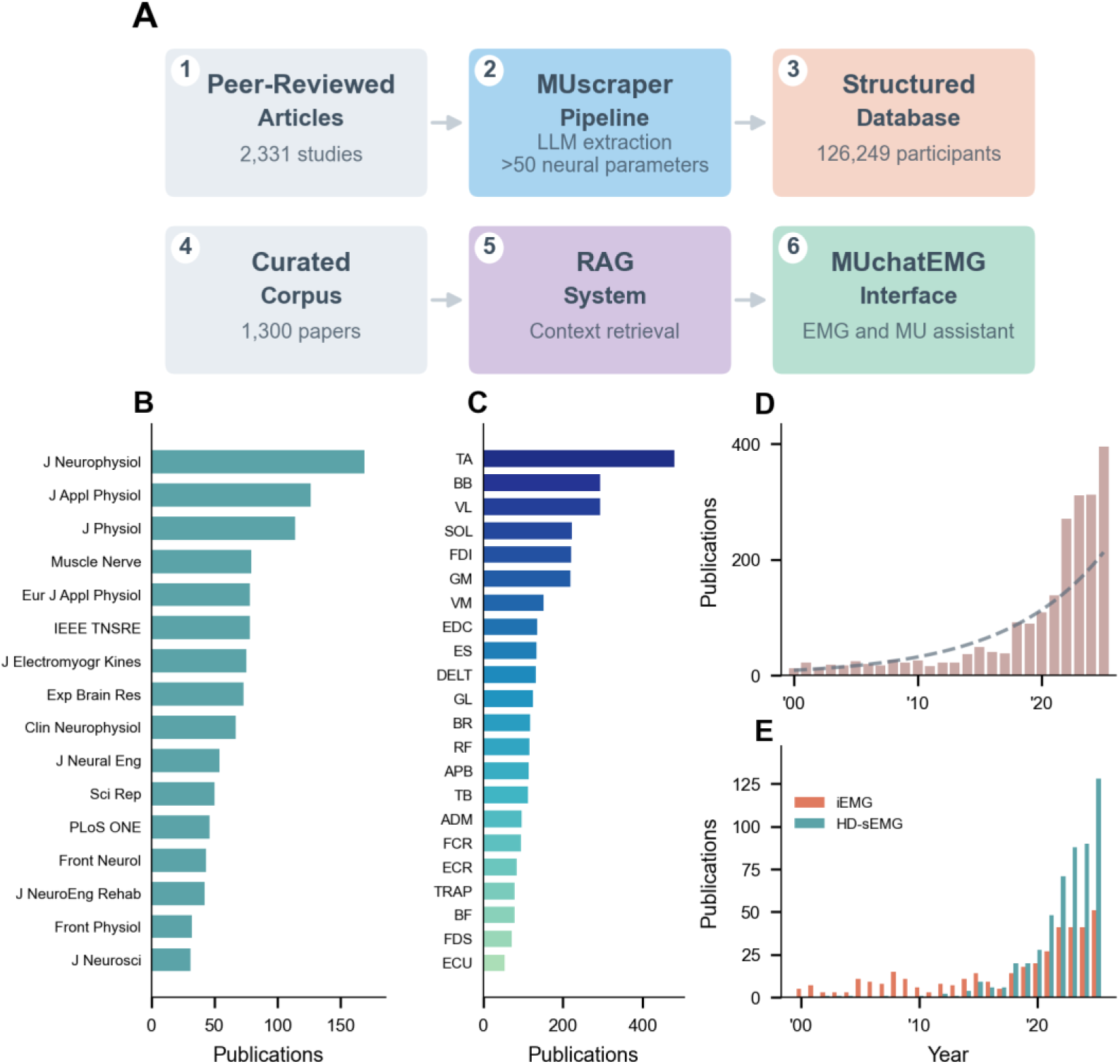
Overview of the NeuromechaniX platform and characteristics of the motor-unit database. (A) Schematic representation of the NeuromechaniX pipeline. The workflow shows peer-reviewed articles on human limb muscles (n=2,331 studies) processed through the MUscraper pipeline, which employs large language model (LLM)-based extraction to retrieve approximately 50 standardized data fields from each publication. The extracted data are compiled into a structured database. The participant count shown (n=30,503 unique participants from 1,774 studies with valid demographic data) reflects reported sample sizes; note that the same individuals may appear in more than one study. A curated corpus of 1,300 full-text papers provides the foundation for a Retrieval-Augmented Generation (RAG) system for the MUchatEMG interface. (B) Distribution of publications across scientific journals. Horizontal bars show the top 17 journals by number of motor-unit publications. The Journal of Neurophysiology leads, followed by Journal of Applied Physiology and Journal of Physiology. (C) Distribution of publications across muscles. Horizontal bar chart indicates the number of studies for each muscle, with tibialis anterior (TA) and biceps brachii (BB) being the most frequently studied. (D) Temporal distribution of motor-unit publications. Bar chart shows the number of studies published per year from 2000 to 2025. The dashed gray line represents a second-order polynomial illustrating exponential growth in motor-unit research. (E) Comparison of intramuscular EMG (iEMG) versus high-density surface EMG (HD-sEMG) publications over time, showing the methodological transition toward HD-sEMG decomposition.

**Table 1.** Sample sizes in NeuromechaniX.

| Component | Studies | Participants | Journals |
| --- | --- | --- | --- |
| Full corpus | 2,331 | 30,503 | 234 |
| Curated corpus (MUchatEMG) | 1,300 | ~18,000 | 187 |
| Motor unit/Force (MUscraper) | 113 | ~2,400 | 48 |

Motor-unit discharge rate analyses required studies to report mean discharge rates with standard deviations, yielding 766 condition-level observations across 113 studies examining seven muscles. The sex-stratified analysis compiled data from the Lulic-Kuryllo and Inglis (2022) systematic review supplemented with studies from 2023-2025, yielding 133 study entries with sex-specific discharge rates. Age-stratified analyses included studies reporting participant age alongside discharge rate or MVC force, yielding 253 and 208 conditions, respectively. Motor-unit counts represent conservative estimates because only ∼41% of conditions reported decomposition yield.

### MUscraper: Automated Metadata Extraction

The MUscraper system addresses the foundational challenge that experimental parameters and study characteristics in the motor-unit literature are reported in heterogeneous narrative formats that challenge manual aggregation. Traditional systematic reviews require human extractors to read each paper and manually transcribe values into spreadsheets, a process that is time-intensive, error-prone, and fundamentally limited in the number of variables that can be captured from each paper. MUscraper automates this extraction process using large language models guided by a comprehensive schema, enabling systematic capture of over 200 structured data fields organized into 17 major sections from each publication.

The extraction schema follows a hierarchical JSON structure aligned with BIDS-BEP042 standards for EMG data (Supplementary S1 provides an exemplary structure), comprising over 200 fields organized into 17 major sections (see Supplementary S2.1 for the complete schema structure). Core sections capture: (1) provenance metadata including schema version and extraction timestamps, bibliographic information including title, DOI, PMID, and author details with ORCID identifiers; (2) participant demographics including sample size, age statistics, sex distribution, health status, and training level; (3) EMG acquisition parameters including electrode type, grid configuration, inter-electrode distance, sampling frequency, and filtering specifications (4) muscles studied with canonical naming following a controlled vocabulary of over 60 muscles organized by anatomical region; (5) task parameters including contraction type, force levels, duration, and joint configuration; (6) decomposition methodology including algorithm family, software, validation method, and quality thresholds; and (7) motor-unit results including discharge-rate statistics, coefficient of variation for interspike interval, recruitment thresholds, and motor-unit action potential characteristics. Extended sections capture specialized measurements including persistent inward currents, common synaptic input, twitch properties, and clinical pathology information when reported.

The extraction pipeline processes each publication through a series of stages designed to maximize accuracy while flagging uncertain extractions for human review. Text is first extracted from PDF documents with section identification, enabling the language model to focus on relevant sections for each field type. We used two different large language models accessed via API: Claude Opus 4.5 (USA) and DeepSeek V3 (China). The complete JSON schema is provided to the language model as an extraction template, with detailed field descriptions and example values guiding the model toward appropriate outputs. The model generates structured JSON conforming to the schema, which is then validated against JSON Schema Draft 2020-12 specifications to ensure type correctness, required field presence, and value constraint satisfaction. Extractions that fail validation or contain values flagged as potentially erroneous based on physiological plausibility checks are routed to manual review.

Terminology harmonization employs a controlled vocabulary (Supplementary S2.2) that resolves synonyms and standardizes units across the heterogeneous reporting conventions found in the literature. Common synonyms such as “firing rate” and “discharge rate” are mapped to canonical terms, muscle abbreviations are expanded to standardized names, and units are converted to consistent formats. This harmonization enables aggregation across studies that may use different terminology for identical concepts. Quality control statistics indicate a schema validation pass rate of 94.2% of extracted records, with common failure modes including missing required fields, type mismatches, and out-of-range values that trigger manual review.

### MUchatEMG: Retrieval-Augmented Generation Architecture

The MUchatEMG system implements a retrieval-augmented text summarization architecture (Lewis et al., 2020) designed for scientific literature synthesis. The system augments the generative capabilities of a language model with retrievable knowledge from the curated corpus, grounding responses in verifiable sources rather than relying solely on parametric knowledge encoded during model training.

The document processing pipeline begins with text extraction from PDF documents using the PyMuPDF library, which preserves structural elements including section boundaries, paragraph breaks, and reading order. The extraction algorithm processes each page sequentially, identifying text blocks through layout analysis while excluding tables and figures that would introduce parsing artifacts. Section boundaries are identified through pattern matching against standard academic headings, and each extracted section is tagged with its type to enable section-aware retrieval during query processing. Documents that fail extraction due to corruption or lack of embedded text are flagged for manual review or optical character recognition processing.

Text segmentation follows a hierarchical chunking strategy that balances retrieval granularity against semantic coherence. Each document is divided into chunks of approximately 256 tokens with 64-token overlap between adjacent chunks to maintain contextual continuity across chunk boundaries. Critically, chunks never span section boundaries, ensuring that each chunk inherits a single section type from its source text. This section awareness proves essential for query routing, as methodological questions benefit from retrieval of Methods sections while quantitative queries are better served by Results content. Each chunk retains rich metadata linking it to the source publication, including DOI, authors, title, journal, year, section type, chunk index, and character offsets within the source document.

The embedding architecture employs the BGE-large-en-v1.5 model (Xiao et al., 2024), which produces 1,024-dimensional dense vector representations optimized for retrieval tasks. This model implements asymmetric encoding, prepending the instruction “Represent this sentence for searching relevant passages:” to all user queries while encoding document chunks without prefix. This asymmetry improves retrieval performance by aligning query representations toward the semantic space of relevant passages. All embeddings are L2-normalized to enable efficient cosine similarity computation via inner product operations.

The retrieval architecture implements a hybrid approach combining dense semantic search with sparse lexical matching, addressing the complementary strengths and weaknesses of each paradigm. Dense retrieval through Facebook AI Similarity Search (FAISS; Johnson et al., 2021) captures semantic similarity, retrieving passages that express related concepts even when vocabulary differs from the query. Sparse retrieval through the BM25 algorithm (Robertson & Zaragoza, 2009) captures exact lexical matches, ensuring that domain-specific terminology and rare technical terms are not overlooked by semantic embeddings alone. Both indices return the top 100 candidate chunks for each query, and results are fused using Reciprocal Rank Fusion with a smoothing constant of 60 and relative weights of 0.7 for dense and 0.3 for sparse results.

Retrieved chunks undergo section-aware score boosting based on classified query intent. The system categorizes queries into four intent types: definition queries seeking conceptual explanations, method queries seeking protocol specifications, factual queries seeking specific quantitative values, and synthesis queries seeking cross-study comparisons. Methods sections receive a 15% score boost for methodological queries, Results sections receive a 10% boost for factual queries, and Abstract sections receive a 5% boost for overview queries. Following score adjustment, the top 50 candidates are reranked using a cross-encoder model (BGE-reranker-large; Xiao et al., 2024) that jointly encodes query-passage pairs to compute fine-grained relevance scores. The top 20 reranked chunks are passed to the language model for answer generation.

Answer generation employs a large language model with temperature set to 0.2 to minimize randomness while preserving fluency. System prompting enforces strict citation-grounding by instructing the model to reference numbered citations for every factual claim, explicitly acknowledge uncertainty when evidence is limited, flag contradictions between sources, and generate explicit “evidence gap” statements when retrieved context is insufficient to answer the query. This citation-grounding approach ensures that every response is traceable to specific source passages, enabling users to verify claims against the original literature.

Response auditability is further enhanced through a verification architecture that generates parallel responses with and without retrieval context. By comparing the retrieval-grounded response against a direct response generated without retrieved context, the system can automatically flag claims unsupported by retrieved evidence. Confidence scores categorize response reliability as high, medium, or low based on retrieval quality metrics and response consistency.

### Statistical Analysis

A critical consideration in meta-analytic synthesis of motor-unit literature is the appropriate unit of analysis. Reported motor-unit counts cannot be pooled across studies because individual motor units are nested within participants, participants within conditions, and conditions within studies, creating a hierarchical data structure where observations are not independent.

To address this challenge while avoiding the pseudoreplication that would result from treating individual motor units as independent observations, we adopted a condition-level analytical approach. Each unique experimental condition, defined by the combination of muscle, task, force level, and study, provides one data point representing the mean discharge rate for that condition. This approach preserves between-study variability as the primary source of variance while enabling valid statistical inference. The discharge-rate analysis included 766 conditions across 113 studies examining seven muscles: biceps brachii (BB), tibialis anterior (TA), first dorsal interosseous (FDI), vastus lateralis (VL), vastus medialis (VM), gastrocnemius (GAS), and soleus (SOL), where k represents the number of studies per muscle.

Given the characteristics of the aggregated data, non-parametric statistical methods were employed. This choice was motivated by several factors: (1) condition-level discharge rate distributions exhibited significant departures from normality as assessed by Shapiro-Wilk tests; (2) sample sizes were unequal across muscles and demographic groups, ranging from k=17 (gastrocnemius) to k=49 (tibialis anterior); (3) Levene’s tests indicated heterogeneous variances across groups; and (4) the relatively small sample sizes in some subgroups (particularly for sex-stratified and age-stratified analyses) precluded reliable estimation of population parameters required for parametric tests. Non-parametric methods provide valid inference under these conditions without requiring distributional assumptions.

Overall differences in discharge rate across muscles were assessed using the Kruskal-Wallis H test, the non-parametric analogue of one-way ANOVA. This omnibus test evaluates whether at least one muscle distribution differs from the others by comparing mean ranks across groups. Following significant omnibus tests, pairwise comparisons were conducted using Mann-Whitney U tests (also known as Wilcoxon rank-sum tests), which test whether observations from one group tend to be larger than observations from another group without assuming equal variances or normal distributions. For the muscle comparison involving 7 groups, this yielded 21 pairwise comparisons, necessitating correction for multiple testing. Bonferroni correction was applied, adjusting the significance threshold to α = 0.05/21 = 0.0024 for pairwise tests.

Effect sizes were computed as Cohen’s d, calculated as the standardized mean difference using pooled standard deviation: d = (M₁ - M₂) / SD_pooled, where SD_pooled = √[((n₁-1)s₁² + (n₂-1)s₂²) / (n₁+n₂-2)]. Effect sizes were interpreted following conventional thresholds: |d| < 0.2 as negligible, 0.2 ≤ |d| < 0.5 as small, 0.5 ≤ |d| < 0.8 as medium, and |d| ≥ 0.8 as large. To quantify uncertainty in effect-size estimates, 95% confidence intervals were computed using non-parametric bootstrap resampling with 1,000 iterations. Each bootstrap iteration sampled with replacement from both groups, computed Cohen’s d, and the 2.5th and 97.5th percentiles of the bootstrap distribution defined the confidence interval bounds. Bootstrap confidence intervals are preferable when distributions are non-normal or sample sizes are small.

Associations between continuous variables (age versus discharge rate, age versus MVC force, force level versus discharge rate) were assessed using both Pearson product-moment correlation coefficients and Spearman rank correlation coefficients. Pearson correlations quantify linear associations and were used when visual inspection of scatterplots suggested approximately linear relations. Spearman correlations, which assess monotonic associations without assuming linearity, were computed as sensitivity analyses. Statistical significance was assessed using two-tailed tests with α = 0.05. For the primary correlation analyses (age-discharge rate, age-MVC, force-discharge rate), no correction for multiple testing was applied as these represented pre-specified hypotheses based on prior literature; however, 95% confidence intervals for correlation coefficients are reported to convey uncertainty.

Sex differences in discharge rate were analyzed using data from the systematic review by Lulic-Kuryllo and Inglis (2022), supplemented with studies published from 2023 to 2025, yielding 85 male and 48 female studies. Overall sex comparisons employed Mann-Whitney U tests with Cohen’s d effect sizes and bootstrap 95% confidence intervals. To examine whether sex differences varied across force levels, data were stratified into low (≤30% MVC), moderate (31-60% MVC), and high (>60% MVC) categories. A permutation test was used to assess the Force × Sex interaction. The observed interaction effect was computed as the difference in sex effects between high and low force levels: Interaction = (Female_high - Male_high) - (Female_low - Male_low). The null distribution was generated by permuting sex labels 5,000 times within each force level, computing the interaction effect for each permutation. The two-tailed p-value was calculated as the proportion of permuted interaction effects with absolute values equal to or exceeding the observed absolute interaction effect. Permutation tests provide exact inference without distributional assumptions and are particularly appropriate when parametric interaction tests would be unreliable due to small or unequal cell sizes.

Age-stratified analyses classified conditions as “Young” when mean participant age was ≤40 years or “Older” when mean participant age was ≥60 years, following the conventions established by Orssatto et al. (2022). Conditions with intermediate ages (40-59 years) were excluded from categorical comparisons to ensure clear separation between groups. This age threshold corresponds approximately to the average onset of age-related motor-unit remodeling documented in longitudinal studies. Between-group differences were assessed using Mann-Whitney U tests with both Welch’s t-test (which accommodates unequal variances) and Mann-Whitney U as complementary approaches; the more conservative p-value was reported. Effect sizes (Cohen’s d) with bootstrap confidence intervals quantified the magnitude of age-related differences.

Distribution-based visualizations were generated using kernel density estimation (KDE) with Gaussian kernels. Bandwidth was set to 0.3-0.4 depending on sample size, selected to balance smoothness against fidelity to the underlying data structure. Values outside the physiological range (4-50 pps for discharge rate, 0-100% for MVC) were excluded prior to density estimation. Ridge plots display vertically stacked density estimates with median indicators, enabling simultaneous comparison across multiple groups. Raincloud plots combine half-violin density estimates, box plots showing median and interquartile range, and individual data points with horizontal jitter to prevent overplotting.

Motor-unit counts were estimated using two methods depending on data availability: direct reporting when papers explicitly stated total motor units decomposed, or calculation as the product of participant count and average number of motor units per participant when both values were reported. Approximately 41% of conditions reported motor-unit yield data; consequently, reported totals represent underestimates of the true motor-unit corpus.

All statistical analyses were performed using Python 3.11 with SciPy 1.11 (scipy.stats module for non-parametric tests, correlation coefficients, and bootstrap procedures), NumPy 1.24 for numerical operations, and Matplotlib 3.7 with Seaborn 0.12 for visualization. Custom functions implemented bootstrap confidence intervals and permutation tests. Statistical significance was defined as p < 0.05 for all tests unless otherwise specified.

### Web Platform

The NeuromechaniX system is deployed as a web application using the Streamlit framework, providing an interactive interface for literature exploration and synthesis. The chat interface enables citation-grounded conversational queries with numbered references linking to source publications. A metadata explorer allows filtering by muscle, year, journal, and decomposition method with export capabilities in CSV and JSON formats. The visualization dashboard provides interactive plots of corpus statistics and extracted metadata distributions. A source viewer enables direct access to the specific passages supporting each citation, ensuring complete transparency and auditability of all generated responses.

## Results

NeuromechaniX comprises two integrated components: MUscraper, an automated metadata extraction pipeline, and MUchatEMG, a citation-grounded retrieval-augmented generation (RAG) interface. Figure 1 shows an overview of the NeuromechaniX platform architecture, depicting the processing pipeline from 2,331 publications through automated metadata extraction to the final structured dataset and interactive web interface.

The corpus spans 234 unique journals, with the Journal of Neurophysiology, Journal of Applied Physiology, and Journal of Physiology contributing the most publications (Fig. 1B). Tibialis anterior and biceps brachii are the most frequently studied muscles (Fig. 1C), reflecting their accessibility for surface EMG recording and importance in both basic research and clinical applications. Publication rates have grown exponentially since 2000, with a notable acceleration after 2015 coinciding with advances in HD-sEMG decomposition algorithms (Fig. 1D). The methodological transition from intramuscular EMG to HD-sEMG is evident in Fig. 1E, with HD-sEMG publications now dominating the field.

### MUscraper: From Unstructured Reports to Structured Metadata

The full corpus encompasses 2,331 peer-reviewed publications on human limb motor units spanning 1925-2025, with the curated RAG corpus containing 1,300 papers across 234 journals. To enable systematic and section-aware retrieval, we processed the full text into section-aware chunks. MUscraper was designed to automate metadata extraction across large corpora. As an initial step, we developed a detailed metadata template (see Supplementary material for the full schema), capturing a wide range of physiological and methodological variables. We then used a large language model to extract unique studies, transforming unstructured text into structured metadata that conformed to the hierarchical schema. The metadata captured includes participant numbers, muscle(s) studied, task and contraction parameters, electrode arrangement, decomposition methods, and motor-unit discharge characteristics.

The analysis of motor-unit discharge characteristics included 766 conditions across 113 studies examining seven muscles, with each condition representing a unique experimental setting (muscle × task × force level). The included studies reported the average and standard deviation of the instantaneous discharge rate. Studies reporting only peak discharge rates from triangular (ramp) contractions without a corresponding mean steady-state rate were excluded; however, studies that reported mean discharge rates during the plateau phase of ramp-and-hold contractions were included. All statistical analyses were performed on condition-level summary statistics (condition means, standard deviations). The number of studies (k) per muscle is indicated in the figure labels.

MUscraper was validated by manual extraction performed by the authors: 20 parameters per paper were independently extracted by hand from 20 randomly selected papers and compared with the automated pipeline output. This analysis resulted in an accuracy of >95% for the automated extraction. The few errors identified were mainly mismatches between DOI registration and publication date, or incomplete motor-unit results for methodology-focused papers where physiological data are not reported in standard formats. We acknowledge that this validation sample (20 papers, ∼1.5% of the corpus) is limited; future work should employ stratified sampling across publication years, methodologies, and muscle groups, with a larger sample (5-10% of the corpus) to assess extraction reliability more comprehensively.

The database reveals several key characteristics of motor-unit research (Figure 2). Most studies employ relatively modest sample sizes (median=13 participants; Fig. 2A). The number of motor units identified per experimental condition varied (median=6, mean=12.6 motor units; Fig. 2B), reflecting the influence of decomposition algorithms, electrode configurations, muscles studied, subcutaneous tissue thickness, muscle architecture, and participant characteristics such as sex and body composition on decomposition yield. The distribution of mean discharge rates (Fig. 2C) centers around 11.5 pps (median), which is located on the steep portion of the sigmoidal relation between discharge rate and applied force (Macefield et al. 1996). The positive correlation between contraction force and discharge rate (r=0.62, p<0.001; Fig. 2E) is consistent with known features of rate coding.

**Figure 2.**
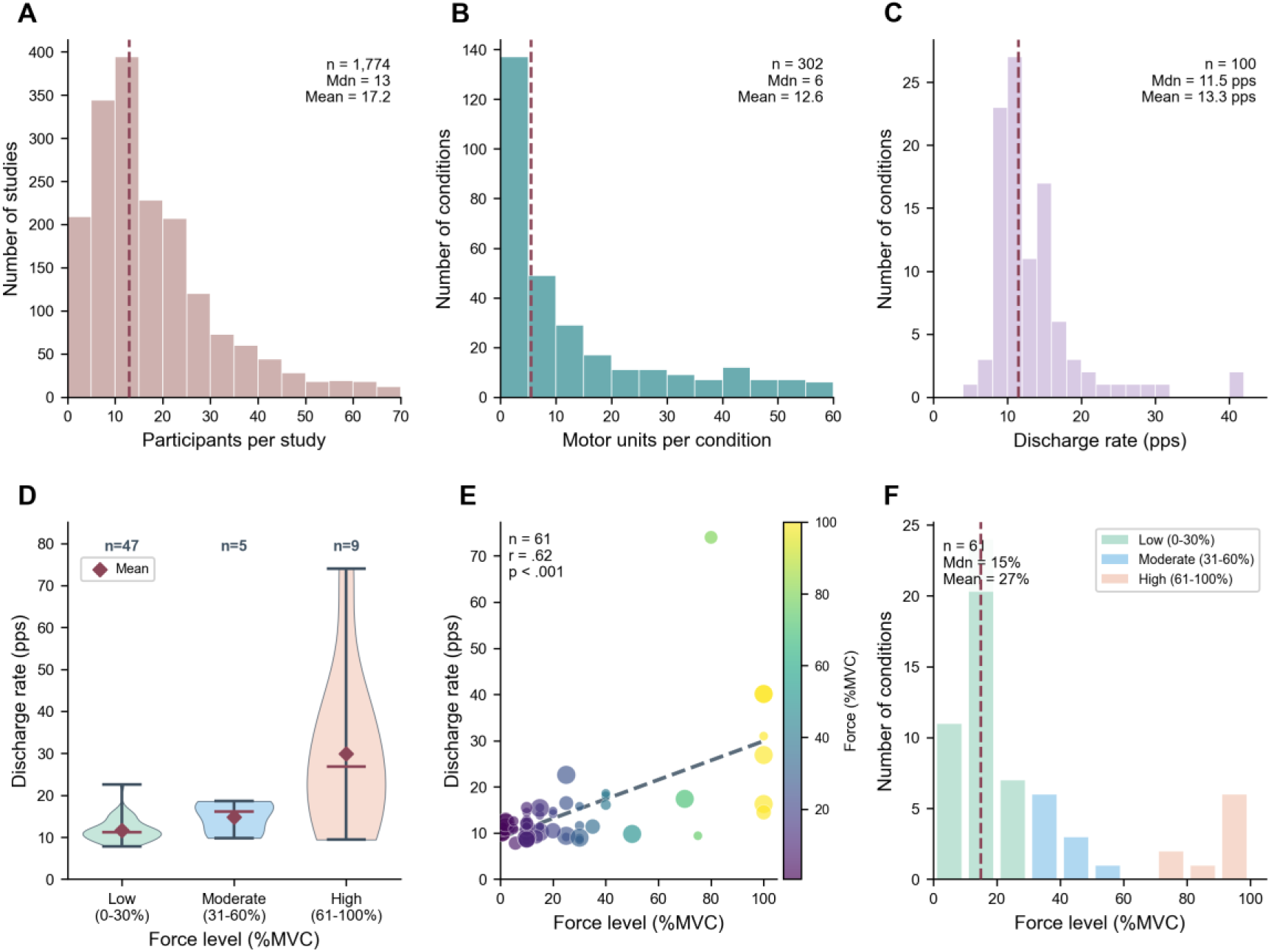
Characteristics of motor-unit studies and the relation between contraction force and motor-unit discharge rate. A condition is defined as a unique combination of muscle, task, and force level within a study. (A) Distribution of participants per study. Histogram shows the number of participants reported in each study (n=1,774 studies with valid participant data). The vertical dashed line indicates the median value (Mdn=13 participants). Statistics: median=13, mean=17.2 participants per study. (B) Distribution of motor units per condition. Histogram plots the number of motor units identified per experimental condition (n=302 conditions with valid motor unit counts). The vertical dashed line indicates the median value (Mdn=6 motor units). Statistics: median=6, mean=12.6 motor units per condition. (C) Histogram indicates the distribution of mean discharge rate reported across experimental conditions (n=100 conditions). The vertical dashed line indicates the median value (Mdn=11.5 pps). Statistics: median=11.5 pps, mean=13.3 pps. (D) Discharge rate by target force. Violin plots show the distribution of mean discharge rates grouped by contraction intensity: Low (0-30% MVC, n=47), Moderate (31-60% MVC, n=5), and High (61-100% MVC, n=9). (E) Relation between force level and discharge rate. Scatter plot of the correlation between contraction intensity (%MVC) and mean motor-unit discharge rate (n=61 conditions). Statistics: Pearson r=0.62, p<0.001. (F) Distribution of force levels tested across experimental conditions (n=61). The vertical dashed line indicates the median (15% MVC); mean=27% MVC.

Even though HDsEMG decomposition can identify motor units reliably up to 80 % maximal voluntary contraction (MVC) force, most studies focus on low forces (median=15% MVC; Fig. 2F), with relatively few studies examining discharge characteristics above 60% MVC force. This finding represents an important gap in the literature. One notable outlier in Fig. 2E reports a discharge rate exceeding 30 pps at high force, likely reflecting peak instantaneous rates that substantially exceed steady-state means.

### Muscle-Specific Discharge Rate Characteristics

Analysis of discharge rates across seven muscles (Figure 3) revealed substantial and statistically significant differences between muscles (Kruskal-Wallis H test, p<0.001). The dataset comprises 766 conditions across 113 studies examining seven muscles: biceps brachii (BB), tibialis anterior (TA), first dorsal interosseous (FDI), vastus lateralis (VL), vastus medialis (VM), gastrocnemius (GAS), and soleus (SOL). These seven muscles were selected because each had enough studies (k≥10) reporting both mean and standard deviation of discharge rate to permit robust statistical comparison. Other muscles in the corpus had fewer studies reporting these specific outcome measures and were excluded from the discharge-rate analysis but are available in the MUscraper database for future analyses.

**Figure 3.**
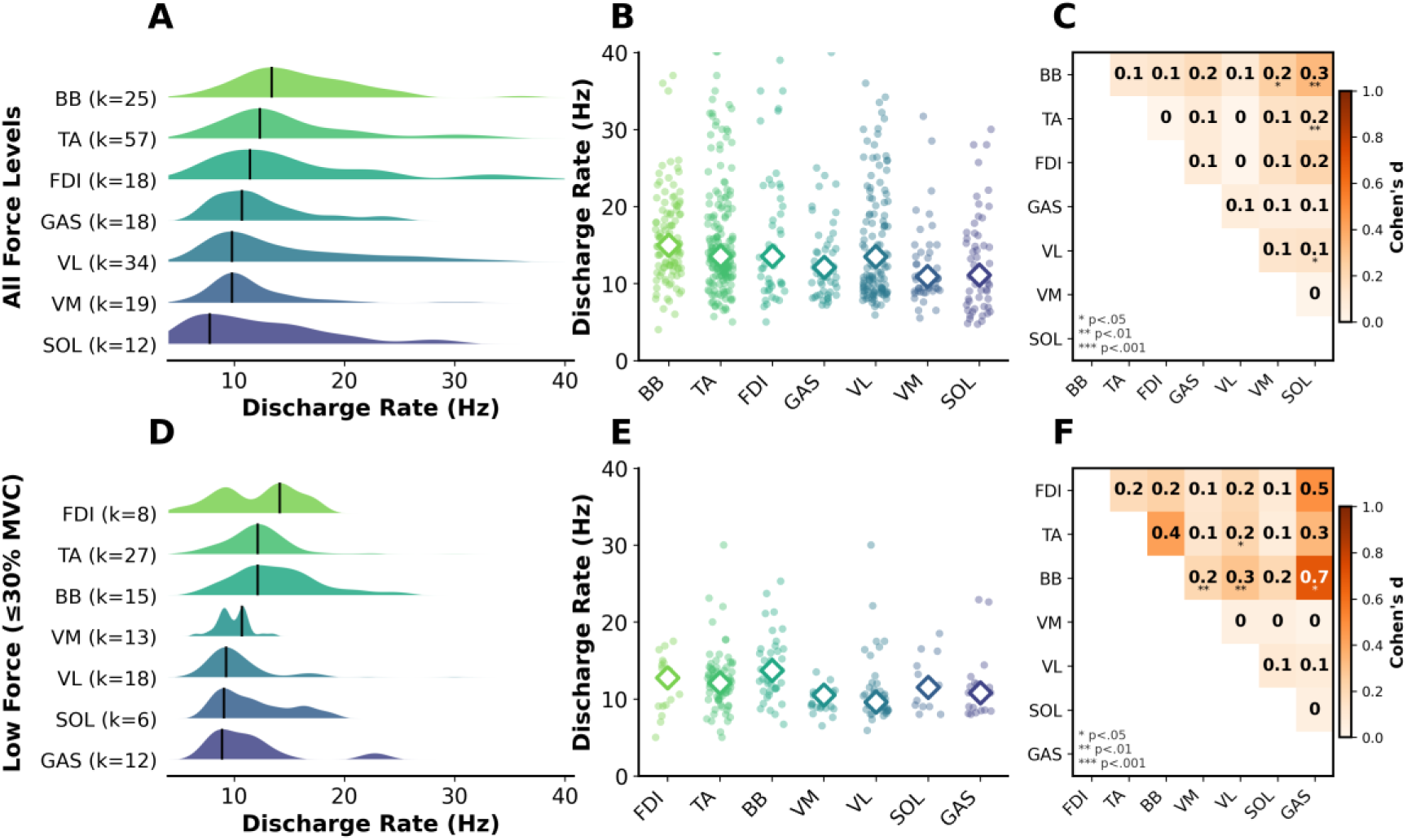
Distributions of motor-unit discharge rate across seven muscles. Top row (A-C): Discharge-rate analysis across all force levels. (A) Ridge plots of discharge-rate distributions by muscle, with muscles ordered by median discharge rate. Each ridge plot represents the kernel density estimate (KDE) of discharge rates for a single muscle. Black vertical lines indicate the median of each distribution. Labels show muscle abbreviation and number of studies (k). (B) Strip plots of discharge rate by muscle. Individual data points represent mean discharge rates for one study condition. White diamond markers indicate the median across all observations for each muscle. (C) Effect-size map for pairwise muscle comparisons. Matrix displays Cohen’s d effect sizes. Statistical significance indicated by asterisks (* p<0.05, ** p<0.01, *** p<0.001). Bottom row (D-F): Discharge-rate analysis restricted to low-force conditions (≤30% MVC). (D-F) Same analyses for studies that used low-force contractions. Abbreviations: BB, Biceps Brachii; TA, Tibialis Anterior; FDI, First Dorsal Interosseous; VL, Vastus Lateralis; VM, Vastus Medialis; GAS, Gastrocnemius; SOL, Soleus.

Ridge-plot distributions sorted by median discharge rate (Fig. 3A) reveal a clear hierarchy: biceps brachii exhibits the highest median discharge rate (15.9 pps), followed by first dorsal interosseous (13.7 pps), tibialis anterior (13.5 pps), vastus lateralis (12.3 pps), gastrocnemius (11.3 pps), vastus medialis (10.7 pps), and soleus (9.9 pps). Importantly, force levels were distributed similarly across muscles (Kruskal-Wallis H=8.2, p=0.22), indicating that the observed hierarchy reflects genuine physiological differences. These findings are derived from an estimated pool of ∼160,000 motor units across the seven muscles, with the high-discharge-rate muscles (biceps brachii, first dorsal interosseous, tibialis anterior; median >13 pps) contributing ∼124,000 motor units, whereas the low-discharge-rate muscles (vastii, gastrocnemius, soleus; median <13 pps) contribute ∼39,000 motor units.

The corresponding strip plots (Fig. 3B) confirm this pattern, displaying individual condition-level observations. Effect sizes (Fig. 3C) were small-to-moderate as indicated by Cohen’s d values (0.0-0.7), reflecting consistent differences in discharge rate between muscles. Fig. 3D-F show the same analysis restricted to low force conditions (≤30% MVC). The muscle ranking persists at low force levels, with biceps brachii, FDI and tibialis anterior showing the highest discharge rates.

### Muscle-Specific Force Characteristics

Beyond discharge rate, muscles differ substantially in force-generating capacity. Analysis of MVC forces across ten muscles (Figure 4)—a larger set than the seven used for discharge-rate analysis because MVC force is more widely reported than discharge-rate statistics—reveals a hierarchy spanning nearly two orders of magnitude, from small intrinsic hand muscles to large thigh extensors. The vastii (vastus lateralis and medialis combined) produced the highest MVC forces during knee-extension tasks, followed by triceps brachii (elbow extensors), brachialis (elbow flexors), and tibialis anterior (dorsiflexors), with first dorsal interosseous (FDI) and abductor pollicis brevis (APB) producing the lowest forces (Fig. 4A). Data were pooled across all tasks to capture the full variability in force; task-specific parameters are preserved in MUscraper for more granular analyses. It should be noted that MVC forces reflect the action performed (e.g., elbow flexion, knee extension) reflecting the synergistic action of multiple muscles.

**Figure 4.**
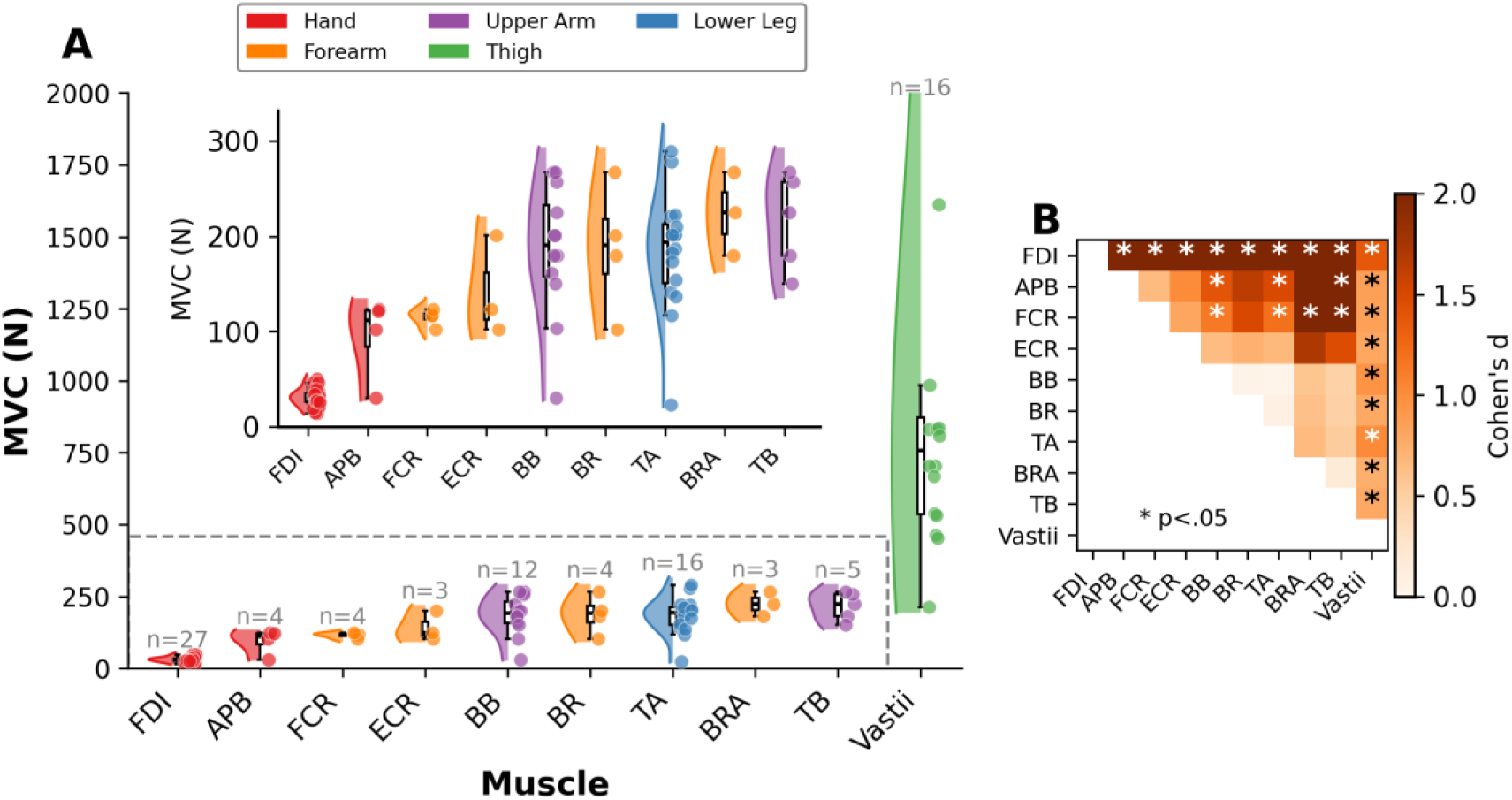
Maximal voluntary contraction (MVC) force across muscles. (A) Raincloud plots of MVC force by muscle. Distribution of MVC forces (Newtons) for 10 muscle groups sorted by median force. Each raincloud comprises: (1) a half-violin plot of the kernel-density estimate on the left, (2) a boxplot shows median and interquartile range in the center, and (3) individual data points on the right, where each point represents the mean MVC force reported for one experimental condition within a single study. Sample sizes (n) are shown above each distribution. Muscles are color-coded by anatomical region: Hand (red), Forearm (orange), Upper Arm (purple), Thigh (green), Lower Leg (blue). An inset panel shows a zoomed view of muscles with lower MVC forces (excluding Vastii). (B) Pairwise effect-size matrix. Upper-triangular map plots Cohen’s d effect sizes for all pairwise muscle comparisons. Color intensity indicates effect-size magnitude (0-2 scale). Asterisks (*) indicate statistically significant differences (Mann-Whitney U test, p<0.05). Abbreviations: FDI, First Dorsal Interosseous; APB, Abductor Pollicis Brevis; FCR, Flexor Carpi Radialis; ECR, Extensor Carpi Radialis; BB, Biceps Brachii; BR, Brachioradialis; TA, Tibialis Anterior; BRA, Brachialis; TB, Triceps Brachii; Vastii, Vastus Lateralis/Medialis.

The apparent overlap in force ranges between anatomically disparate muscles (e.g., APB and BB) reflects differences in transducer setups, moment arms, and the specific actions measured across studies rather than comparable force-generating capacity; for example, BB force is typically measured during elbow flexion with a moment arm of ∼30 cm, whereas APB force is measured during thumb abduction with a much shorter moment arm, and the force instead of the torque being reported.

The raincloud visualization (Fig. 4A) displays the full distribution of MVC values for each muscle, with muscles sorted by median force and color-coded by anatomical region: hand muscles (red), forearm muscles (orange), upper arm muscles (purple), thigh muscles (green), and lower leg muscles (blue). The inset panel provides a zoomed view of the smaller muscles (excluding vastii) for clearer visualization of their distributions.

Effect-size analysis (Fig. 4B) demonstrates large Cohen’s d values (>0.8) for comparisons between muscles from different anatomical regions, particularly between hand muscles and proximal limb muscles.

### Sex Differences in Motor-Unit Discharge Rate

Sex differences in motor-unit discharge rates were analyzed using data compiled from the literature, including the systematic review by Lulic-Kuryllo and Inglis (2022) and additional studies from 2023-2025 (Figure 5). The analysis included 133 studies: 85 male and 48 female entries. Overall, females exhibited higher discharge rates than males (median 14.5 vs 11.9 pps; mean 15.6 vs 13.3 pps), a difference of approximately 2.5 pps or 20% relative to male values (Mann-Whitney U=1532, p=0.018, Cohen’s d=0.38; Fig. 5B). Although effect size is typically classified as “small,” the absolute difference is comparable to the increase in discharge rate observed when force is doubled from 10% to 20% MVC, suggesting physiological relevance.

**Figure 5.**
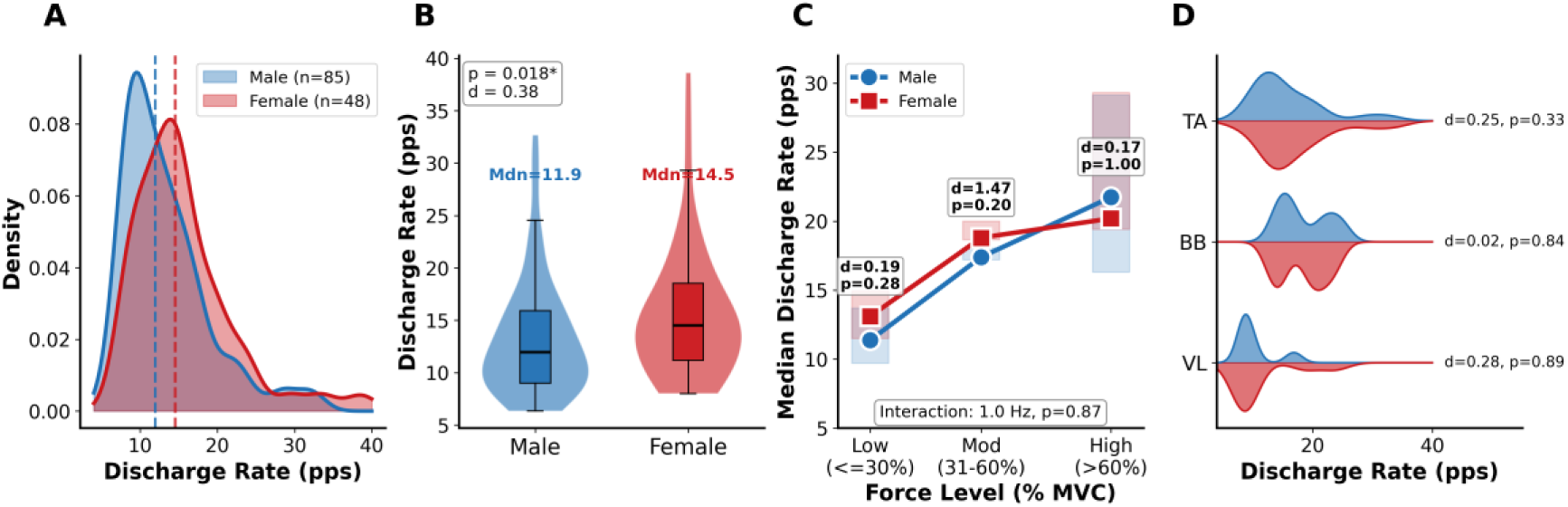
Analysis of sex differences in motor-unit discharge rates. (A) Overall distribution of discharge rates by sex. Kernel-density estimation (KDE) plots of the distribution of mean discharge rates for males (blue, n=85) and females (pink, n=48). There was substantial overlap in the distributions, with females showing a slight rightward shift toward higher discharge rates. (B) Sex comparison of discharge rates. Violin plots comparing mean discharge rates between males and females. The box within each violin plot shows the interquartile range (IQR) with the median indicated by the horizontal line. Statistical comparison: Mann-Whitney U=1532, p=0.018, Cohen’s d=0.38 (small effect). Discharge rates were significantly greater for females (median 14.5 pps) than males (median 11.9 pps). (C) Force level × Sex interaction. Line plot showing mean discharge rates across three force categories (Low: ≤30% MVC, Moderate: 31-60% MVC, High: >60% MVC) for males (blue) and females (pink). Error bars represent ±1 SEM. Permutation test for interaction: difference=1.0 pps, p=0.87 (not significant). (D) Muscle-specific sex comparisons. KDE distributions showing discharge rates by sex for three muscles with sufficient data (n≥3 per sex): Tibialis Anterior (TA), Biceps Brachii (BB), and Vastus Lateralis (VL). Data source: Compiled from Lulic et al. (2022) systematic review tables plus new studies from 2023-2025.

Examination of the influence of sex on the relation between discharge rate and force (Fig. 5C) revealed no significant force × sex interaction (permutation test: p=0.87), indicating that the sex difference in discharge rate is maintained across low, moderate, and high forces. However, the number of studies available for such comparisons is limited (see below and Discussion section).

Muscle-specific analysis (Fig. 5D) showed that tibialis anterior, biceps brachii, and vastus lateralis had sufficient data for sex comparisons (n≥3 per sex). Although females tended to show higher discharge rates across these muscles, the differences were not statistically significant when analyzed separately (TA: p=0.33, d=0.25; BB: p=0.84, d=0.02; VL: p=0.89, d=0.28), likely due to the low sample sizes when stratified by muscle.

The larger male sample size reflects a broader pattern of sex imbalance in the motor unit literature. This imbalance limits the statistical power for sex-stratified analyses and underscores the need for more balanced sex representation in future studies.

### Age-Related Changes in Motor-Unit Properties

The motor-unit literature is dominated by studies on young adults. Analysis of publication distribution by participant age (Fig. 6A) reveals that 1,516 publications (58%) studied participants under 30 years of age, whereas only 399 publications (15%) examined adults over 60 years. This age imbalance reflects both the convenience of recruiting university students and the challenges associated with recruiting older individuals.

**Figure 6.**
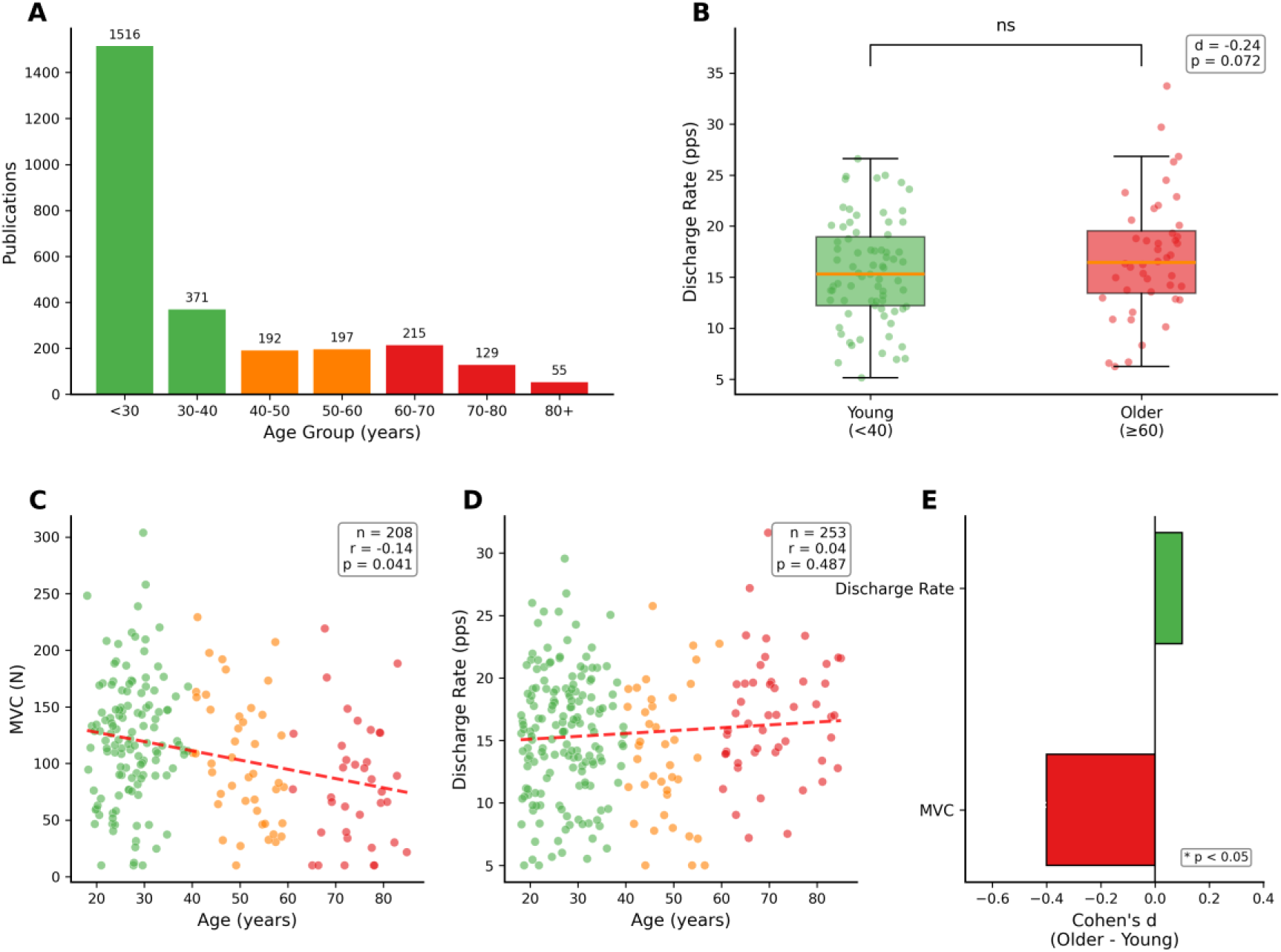
Analysis of age-related changes in motor-unit discharge rates and maximal voluntary contraction force. (A) Publication distribution by age group. Bar chart showing the number of publications stratified by mean participant age. Most motor-unit studies involved young adults (<30 years, n=1,516), with fewer studies on older adults (<40: n=371; 40-50: n=192; 50-60: n=197; 60-70: n=215; 70-80: n=129; 80+: n=55). (B) Discharge-rate comparison: Young vs. Older. Box plots with overlaid data points comparing mean discharge rates between young (<40 years, green) and older (≥60 years, orange) adults. Statistical comparison: Cohen’s d=-0.24, p=0.072 (not significant). The distributions overlap substantially, suggesting no detectable age difference in discharge rate when pooling across studies. (C) Relation between age and MVC force. Scatter plot showing the correlation between participant age and MVC force (n=208 conditions). Statistics: Pearson r=-0.14, p=0.041, indicating a significant age-related decline in MVC force. Point colors indicate age category (green=young, orange=middle-aged, red=older). (D) Relation between age and mean motor-unit discharge rate was not significant (n=253 conditions). Statistics: Pearson r=0.04, p=0.487. (E) Effect-size summary. Bar plot of Cohen’s d effect sizes for age-related differences. MVC force shows a significant negative effect (older < young, *p<0.05), whereas discharge rate shows a negligible, non-significant effect.

Despite the established view that motor-unit discharge rates decline with aging, our cross-study analysis reveals a more nuanced conclusion. Comparison of discharge rates between young (<40 years) and older (≥60 years) adults (Fig. 6B) found no statistically significant difference (Cohen’s d=-0.24, p=0.072). The distributions overlap substantially, with both groups exhibiting median discharge rates near 15 pps.

Although discharge rate was not significantly correlated with age (Fig. 6D; n=253, r=0.04, p=0.487), there was a significant negative correlation with MVC force (Fig. 6C; n=208, r=-0.14, p=0.041). The effect-size summary (Fig. 6E) illustrates this dissociation: MVC force declines with aging (d≈-0.4, p<0.05), whereas discharge rate shows a negligible, non-significant effect (d≈0.1). Both findings underscore the limited representation of middle-aged and older adults in the motor-unit literature, with only 27% of publications examining participants over 40 years of age. Critically, the categorization of cohorts based on chronological age overlooks the presence of low and high performers across the lifespan (Ingram et al., 2023; Lazarus et al., 2019; Pollock et al., 2015; Wunderle et al., 2024).

### Corpus-Level Text Analysis

Text analysis of the curated corpus (1,300 full-text papers) reveals the semantic structure of the motor-unit literature. The word cloud (Fig. 7) displays the most frequent domain-specific terms, with “motor”, “unit”, “muscle”, “force”, and “discharge rate” dominating, representing the core concepts of motor-unit physiology.

**Figure 7.**
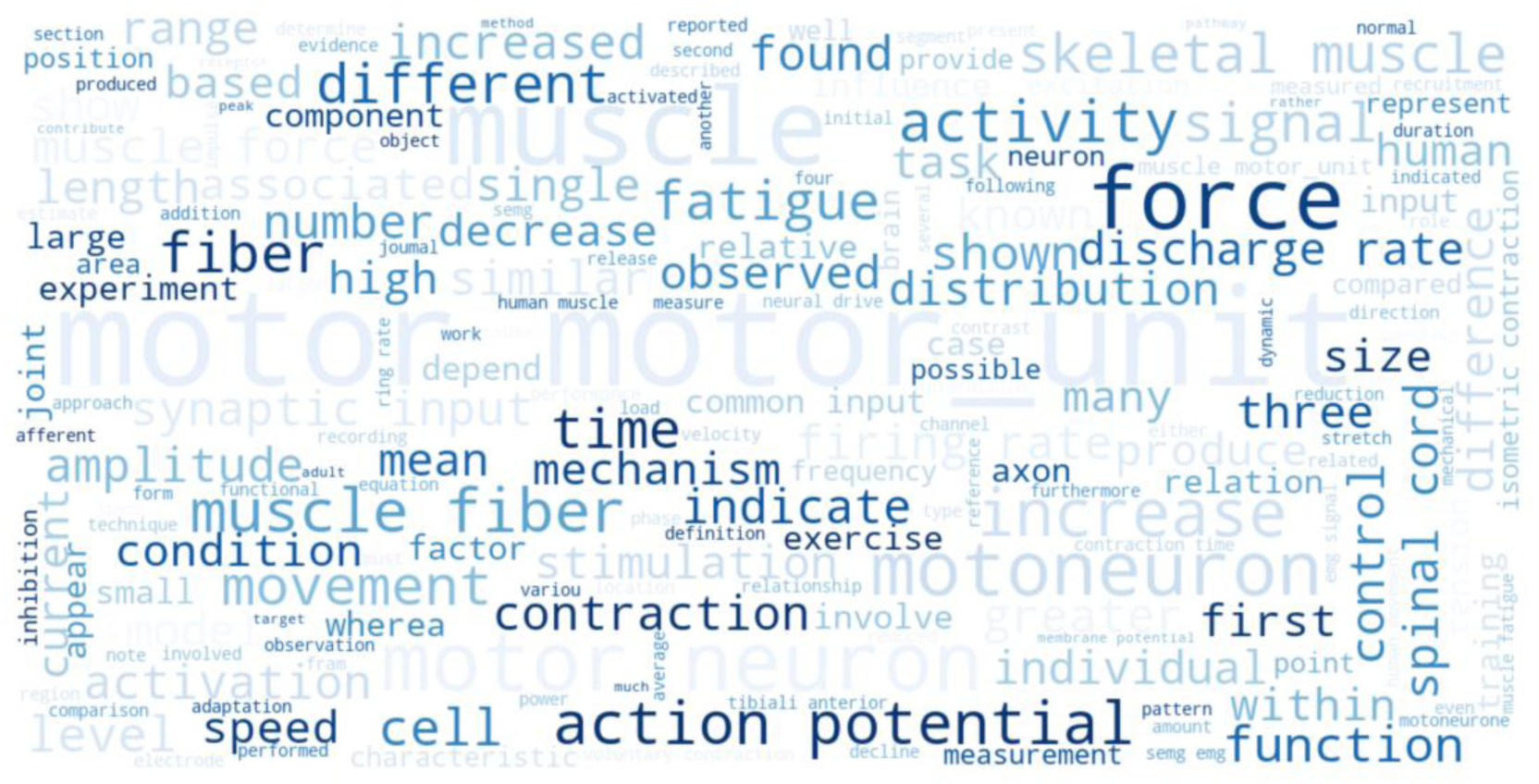
**Word cloud of the NeuromechaniX corpus**. Word cloud showing the most frequent terms across the curated corpus of 1,300 motor unit and EMG publications. Word size reflects term frequency. Dominant terms include “motor”, “unit”, “muscle”, “force”, “discharge rate”, “activity”, and “action potential”, representing the core concepts of motor unit physiology.

The word usage shift analysis (Fig. 8) reveals a striking temporal transition in scientific vocabulary around the year 2000: before 2000, the literature included “tension”, “motoneurone” (British spelling), “needle”, and “tetanic”—vocabulary characteristic of classical neurophysiology and intramuscular recording. Since 2000, there is increased use of “participant”, “decomposition”, “coherence”, and “corticospinal”—reflecting the shift toward HD-sEMG decomposition and systems-neuroscience approaches. This change in vocabulary mirrors the methodological transition documented in Fig. 1E. The literature is unevenly distributed across muscles (vastus lateralis, tibialis anterior, and biceps brachii most studied) and shows substantial gaps in demographic reporting (∼90% male), limiting sex-stratified syntheses.

**Figure 8.**
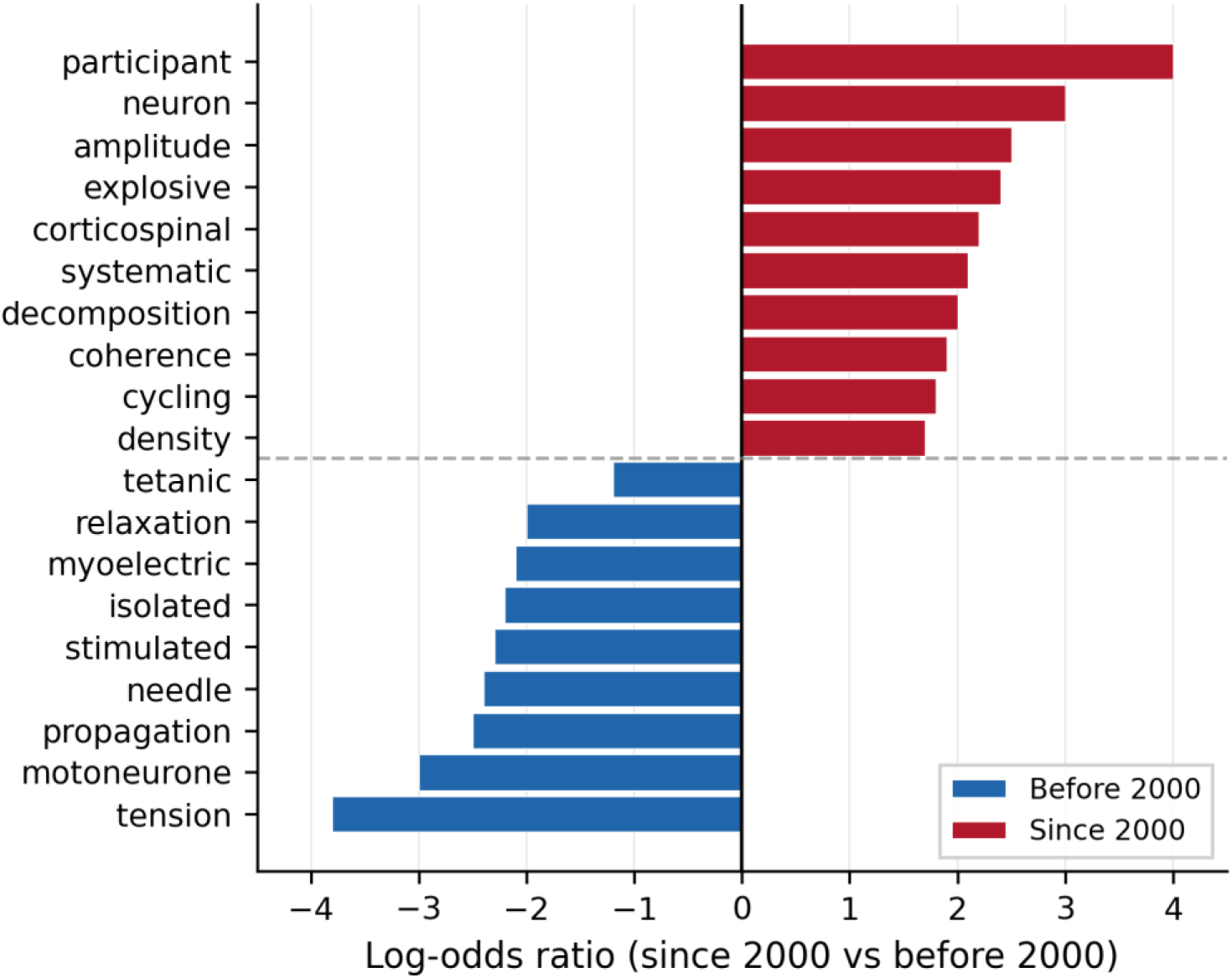
Word usage shift around the year 2000. Horizontal bar chart showing log-odds ratios for terms that differ significantly in frequency between before and after 2000. Blue bars indicate terms more frequent in the earlier literature (classical neurophysiology terminology: “tension”, “motoneurone”, “needle”, “stimulated”, “tetanic”), whereas the red bars denote terms more frequent since 2000 (modern terminology: “participant”, “neuron”, “decomposition”, “coherence”, “corticospinal”). The dashed horizontal line separates the two temporal categories. This vocabulary shift reflects the transition from classical intramuscular techniques to HD-sEMG decomposition and systems-neuroscience approaches.

### MUchatEMG: Supplementary Text Summarization Interface

We additionally provide MUchatEMG (https://neuro-mechanix.com/), a web-based text summarization interface that retrieves and summarizes passages from the curated corpus with citation anchoring. MUchatEMG is not designed as an alternative to general-purpose language models; rather, it constrains responses to the peer-reviewed corpus, ensuring that every factual claim is traceable to specific publications.

All responses include numbered citations linking to source metadata (DOI, authors, title, journal, year). MUchatEMG returns structured responses organized into five tiers: grounded facts with direct citations, synthesized cross-study findings, identified evidence gaps, extended reasoning, and testable hypotheses (see Supplementary Tables S3-S4 for representative examples). Technical details of the retrieval architecture are provided in the Methods.

### Open-Access Metadata Explorer

To enable community access to the structured database, we provide an open-access interactive web platform (https://neuro-mechanix.com/metadata). The metadata explorer (Fig. 9) allows researchers to filter studies by muscle, toggle the visibility of extracted metadata fields across multiple categories (core bibliographic information, motor-unit variables, neural interfacing parameters, and hardware/protocol specifications), search across the entire database, and export filtered datasets in tabular format. This interface exposes the full breadth of the over 200 structured fields extracted by MUscraper, enabling researchers to perform custom queries and retrieve study-level metadata without requiring programming expertise. The platform is designed to be community-extensible, with plans to allow certified researchers to contribute publications to the curated corpus.

**Figure 9.**
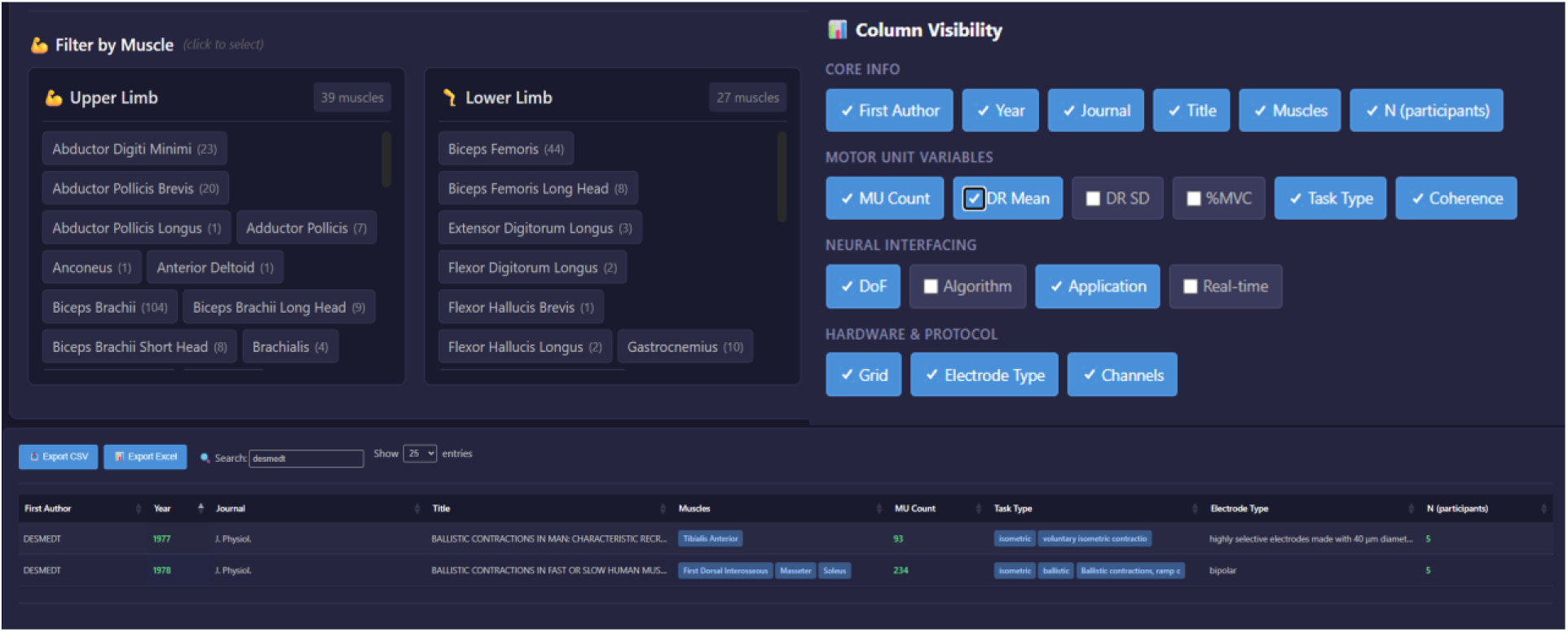
The NeuromechaniX open-access metadata explorer. Screenshot of the interactive web platform (https://neuro-mechanix.com/metadata) for exploring the structured motor-unit database. The interface comprises three main components: (left) a muscle filter panel organized by anatomical region (upper limb and lower limb), allowing users to select specific muscles and view the number of available studies per muscle; (right) a column visibility panel that allows users to toggle the display of extracted metadata fields organized into four categories—Core Info (first author, year, journal, title, muscles, sample size), Motor-Unit Variables (motor-unit count, discharge rate mean and standard deviation, motor-unit conduction velocity, task type, coherence), Neural Interfacing (DOI, algorithm, application, real-time capability), and Hardware/Protocol (grid configuration, electrode type, number of channels); (bottom) a searchable, sortable data table displaying the filtered study-level metadata with pagination and export functionality. The platform enables researchers to explore, filter, and download the complete structured database extracted by MUscraper without requiring programming expertise.

## Discussion

This work introduces NeuromechaniX, a domain-specific platform for automated extraction and meta-analysis of the motor-unit literature. The primary contributions are: (1) MUscraper, an automated extraction pipeline that transforms thousands of unstructured publications into structured metadata across over 200 fields organized into 17 major sections, and (2) meta-research insights enabled by the emergent database, including the largest cross-study analysis of motor-unit discharge rates to date. By transforming unstructured reports into structured metadata, NeuromechaniX enables cumulative inference across corpora that exceed the capacity of manual review.

### Automated extraction as scientific infrastructure

MUscraper addresses a fundamental bottleneck in motor-unit meta-research: the inability to programmatically aggregate experimental parameters reported in heterogeneous narrative formats across thousands of publications. Traditional systematic reviews can realistically capture a few variables from each paper; MUscraper captures over 200 structured fields per publication, enabling multi-dimensional analyses that cross-tabulate muscles, tasks, force levels, demographics, and methodological parameters simultaneously.

In practice, MUscraper supports both everyday neuroscience workflow: finding methods, identifying task parameters, locating canonical evidence) and higher-level meta-research (mapping which muscles, tasks, and populations dominate the literature. The structured database enables hypothesis generation through comparative analysis, exposing underexplored connections across decades of work.

### Gaps in muscle representation

The motor-unit literature contains substantial gaps in muscle representation. Postural muscles like soleus and gastrocnemius are underrepresented despite their functional importance for balance and locomotion. Even more striking is the limited data on trunk and core muscles (erector spinae, multifidus, abdominal muscles), shoulder muscles (deltoid, rotator cuff), and respiratory muscles (diaphragm, intercostals). Notably, the current corpus does not yet incorporate the extensive body of work on single-unit firing properties of respiratory muscles (diaphragm, intercostals) obtained using intramuscular electrodes. The current focus on limb muscles studied primarily with HD-sEMG decomposition means that important motor-unit data obtained through intramuscular techniques in these muscle groups remain to be integrated.

These gaps reflect both technical challenges (deep muscle access, movement artifacts) and historical research priorities, but they limit our understanding of neural control across the full operating range of the motor system.

### Muscle-specific discharge characteristics: Physiological implications

Our analysis of discharge rates across 766 conditions from 113 studies on seven muscles reveals robust muscle-specific differences across force levels and experimental protocols. The finding that biceps brachii exhibits the highest discharge rates (median 15.9 pps), followed by first dorsal interosseous (13.7 pps) and tibialis anterior (13.5 pps), whereas postural muscles (soleus 9.9 pps, vastus medialis 10.7 pps, gastrocnemius 11.3 pps) show lower rates, suggests a fundamental distinction between muscles with high or low discharge rates.

This discharge-rate hierarchy may reflect an organizing principle in motor control: muscles requiring rapid force modulation and precise temporal control (upper limb, distal muscles) operate at higher discharge rates, whereas muscles used for postural control (lower limb, proximal muscles) operate at lower rates. This pattern aligns with known differences in functional demands. Upper limb muscles involved in skilled manipulation (biceps brachii, first dorsal interosseous) require rapid force modulation and precise control, which may be facilitated by higher discharge rates and greater rate coding. In contrast, postural muscles seem to be optimized for sustained, low-force contractions during standing and walking, consistent with lower discharge rates and greater reliance on slower contracting motor units. Importantly, force levels were distributed similarly across muscles (p=0.22), confirming that the observed hierarchy reflects genuine physiological differences rather than being confounded by contraction intensity. An intriguing observation to emerge from these data is that tibialis anterior, despite being a lower limb muscle, exhibits discharge rates comparable to upper limb muscles (13.5 pps), whereas the vastii muscles discharge at substantially lower rates (10.7-12.3 pps).

Although the data on MVC force were pooled across all conditions, task-specific parameters are preserved in MUscraper database (e.g., isometric knee flexion/extension, index finger abduction, lengthening and shortening), enabling future studies to examine force-task relations in greater detail.

The substantial variability in motor-unit yield across muscles (median range: 4.3-19.5 MU/participant) has important methodological implications. The high yield in tibialis anterior likely reflects its superficial location and favorable architecture for HD-sEMG recording, whereas deep muscles like trapezius produce lower yields.

### Sex differences and Aging: The need for balanced representation

The marginal but significant sex difference in discharge rate (Cohen’s d=0.38) observed in our analysis adds to growing evidence that motor-unit activity likely differs between males and females. However, the most striking finding from our sex analysis is the dramatic imbalance in the literature: ∼90% of observations come from male participants. This imbalance limits our ability to draw robust conclusions about sex differences and highlights an urgent need for more balanced sex representation in motor-unit research. Given potential sex differences in fiber-type proportions, motor-unit size, and hormonal influences on neuromuscular function, this gap represents a concern. An additional technical consideration is that HD-sEMG decomposition may be subject to sex-specific detection bias: greater subcutaneous adipose tissue thickness in females attenuates surface potentials and may result in preferential detection of larger, more superficial motor units, potentially biasing the reported discharge rate distributions. This detection asymmetry could contribute to the observed sex difference in discharge rate and should be considered when interpreting sex-stratified results from HD-sEMG studies.

Our age-stratified analysis yielded a surprising null result: no significant difference in discharge rate between young (<40 years) and older (≥60 years) adults (d=-0.24, p=0.072), despite significant age-related decline in MVC force (r=-0.14, p=0.041). This finding stands in apparent contradiction to the well-powered meta-analysis by Orssatto et al. (2022), which concluded that motor-unit discharge rates are significantly lower in older adults (SMD=0.66, p=0.001) with effects magnified at higher contraction intensities.

This discrepancy illustrates a fundamental methodological issue in meta-research. Orssatto et al. analyzed 29 studies that compared young and older adults within the same laboratory, using identical equipment, protocols, and investigators. This within-study design controls for the substantial between-laboratory variability in motor-unit recording and decomposition. In contrast, our analysis pools data across independent studies, each of which may have studied young or older participants under laboratory-specific conditions. The methodological heterogeneity between laboratories (electrode configurations, decomposition algorithms, force-normalization procedures, muscle selection) introduces variance that may overwhelm the relatively modest aging effect on discharge rate. One likely explanation for this finding is that the current literature lacks sufficient sample sizes of older adults to detect moderate effects in cross-laboratory analyses. Alternatively, the small age effect may be attributable to the lack of control for the heterogeneity in performance capabilities among cohorts defined by chronological age.

Several specific factors likely contribute to the attenuation of detectable age effects in cross-study pooling. First, Orssatto et al. found that age differences scale with contraction intensity, increasing by 0.009 SMD per 1% MVC. Our database is dominated by low-force studies (median 15% MVC), where the effects of age are smallest. Second, Orssatto et al. found clear age differences in tibialis anterior (SMD=0.78) but not first dorsal interosseous (SMD=0.04), suggesting that intrinsic hand muscles may be protected from age-related discharge rate decline. Third, studies finding expected age differences may be more likely to report and publish age-stratified results, whereas studies finding null results may collapse across age groups without comment. Despite these limitations, cross-laboratory analyses remain valuable precisely because they capture the real-world variability in motor-unit research, provided the cohorts are categorized based on relevant morphometric factors or performance on functional motor assessments.

### The hidden dimension: Why muscle-specific sex and age effects matter for understanding neural control

Perhaps the most important finding from our sex and age analyses is not what we found, but what we could not determine: muscle-specific differences. When stratifying sex comparisons by muscle, only three muscles (tibialis anterior, biceps brachii, vastus lateralis) had sufficient sample sizes (n≥3 per sex) to permit even rudimentary statistical comparison, and none revealed significant sex differences (all p>0.3). The overall sex difference we observed (females higher discharge rate, d=0.38) could reflect a genuine biological phenomenon operating uniformly across the motor system, or it could represent an artifact of muscle sampling bias in the underlying literature. The three muscles we could analyze showed non-significant trends in the same direction (female > male), but with effect sizes ranging from d=0.02 (biceps brachii) to d=0.28 (vastus lateralis). Whether this variation reflects genuine muscle-specific sex differences or sampling noise cannot be determined with current data.

The interaction between sex and age is uncertain. If males and females experience different aging trajectories for discharge rate, and if these trajectories differ across muscles, then pooling across sex and muscle, as most large-scale analyses must do, could obscure genuine effects. The current literature cannot rule out masked interactions.

### Future research focus driven by NeuromechaniX findings

The path forward requires deliberate, targeted data collection. Our analysis identifies specific gaps that should be prioritized: hand muscles in older adults, lower limb muscles in females, and direct within-study sex and age comparisons for the same muscles using identical protocols. The dramatic sex imbalance in the current literature (90% male) makes muscle-specific sex comparisons nearly impossible for most muscles. The concentration of aging studies in a few muscles, combined with methodological heterogeneity across laboratories, limits the power to detect muscle-specific aging effects. The field needs to fill these gaps, perhaps by performing multi-site studies designed to collect sex-balanced, age-stratified data across a standardized muscle battery using harmonized acquisition and decomposition protocols.

Until such data are available, the overall effects we report, whether significant or null, must be interpreted with caution. The significant sex difference (d=0.38) may be real and uniform, or it may be driven by a subset of muscles whereas others show no effect or even opposite effects. The null age effect (d=-0.24) likely reflects a failure of the field to categorize cohorts with appropriate criteria (not chronological age). The NeuromechaniX platform provides the infrastructure to revisit these questions as the evidence base grows. The current results should be used as a quantitative map that indicated where the evidence is sufficient, where it is suggestive, and where it remains fundamentally inadequate to address questions of basic and clinical importance.

### What the current evidence base emphasizes

Across a century of publications, the evidence base is concentrated in a relatively small set of muscles and protocols. Tibialis anterior, biceps brachii, and vastus lateralis account for a large fraction of studies, whereas many trunk and head/neck muscles are under-represented.

The corpus also documents the field’s methodological transition toward widespread HD-sEMG decomposition. These shifts matter for neuroscience interpretation because acquisition and decomposition choices influence which motor unit populations are detected, how discharge characteristics are estimated, and what uncertainty is carried into downstream comparisons.

## Limitations

Several limitations should be acknowledged. First, the atlas reflects the current curated corpus (2,331 publications, 1,300 in RAG) and is therefore incomplete relative to the entire literature. However, this work represents the first systematic extraction and analysis of structured metadata from such a large body of motor-unit research, far exceeding the scope of traditional meta-analyses based on a few variables from dozens of papers.

Second, automated extraction includes schema-based validation and consistency checks but still depends on the clarity and completeness of reporting in the source papers; ambiguous text, non-standard terminology, and complex experimental designs can yield missing or uncertain values that require manual review.

Third, motor-unit counts are conservative estimates because only approximately 40% of experimental conditions explicitly reported decomposition yield; consequently, our totals likely underestimate the actual number of motor units studied across the literature.

Fourth, the current pipeline focuses on text extraction and does not yet incorporate data from figures, tables, or equations, which often contain critical quantitative details not repeated in the narrative text.

Fifth, the dramatic sex imbalance in the source literature (∼90% male) and scarcity of studies in older adults limits the robustness of sex- and age-stratified conclusions.

Sixth, the pooled analyses aggregate data from studies employing heterogeneous experimental protocols, including differences in contraction type (isometric, isotonic, concentric, eccentric), contraction duration, joint angle, rest periods, and the presence or absence of fatigue. These methodological variations are known to influence motor-unit discharge properties and may introduce systematic bias into pooled estimates. Although MUscraper preserves task-specific parameters that enable future subgroup analyses by contraction characteristics, the current analyses pool across these conditions to maximize statistical power. Results should therefore be interpreted as reflecting the central tendency across the published literature rather than controlled experimental comparisons.

## Conclusion

By organizing a century of motor-unit and EMG research into a structured, queryable database, we provide infrastructure for more rigorous synthesis in motor-unit physiology. MUscraper enables automated extraction of over 200 structured metadata fields from thousands of publications, making cross-study statistical comparisons feasible at a scale that manual review cannot achieve. The analysis of discharge rates across hundreds of conditions and studies examining seven muscles reveals robust muscle-specific discharge characteristics and persistent gaps in sex and age representation. This resource is expected to help the community identify evidence gaps, standardize reporting conventions, generate testable hypotheses from cross-study patterns, and accelerate hypothesis-driven research grounded in the published record. As the corpus expands, NeuromechaniX will evolve into an increasingly comprehensive knowledge infrastructure for the neuromechanics field.

## Abbreviations

APB: abductor pollicis brevis
BB: biceps brachii
BR: brachioradialis
BRA: brachialis
ECR: extensor carpi radialis
EMG: electromyography
FCR: flexor carpi radialis
FDI: first dorsal interosseous
GAS: gastrocnemius
HD-sEMG: high-density surface electromyography
iEMG: intramuscular electromyography
MU: motor unit
MVC: maximal voluntary contraction
pps: pulses per second
SOL: soleus
TA: tibialis anterior
TB: triceps brachii
VL: vastus lateralis
VM: vastus medialis.

## Supplementary Information

This supplementary material provides detailed technical documentation for the NeuromechaniX platform, including the complete metadata extraction schema (S1) and controlled vocabulary (S2) used by MUscraper.

### S1: MUscraper JSON Schema (Exemplary Structure)

The MUscraper extraction pipeline uses a hierarchical JSON schema containing over 50 fields organized into seven main sections. Below is a simplified example showing the key structure:

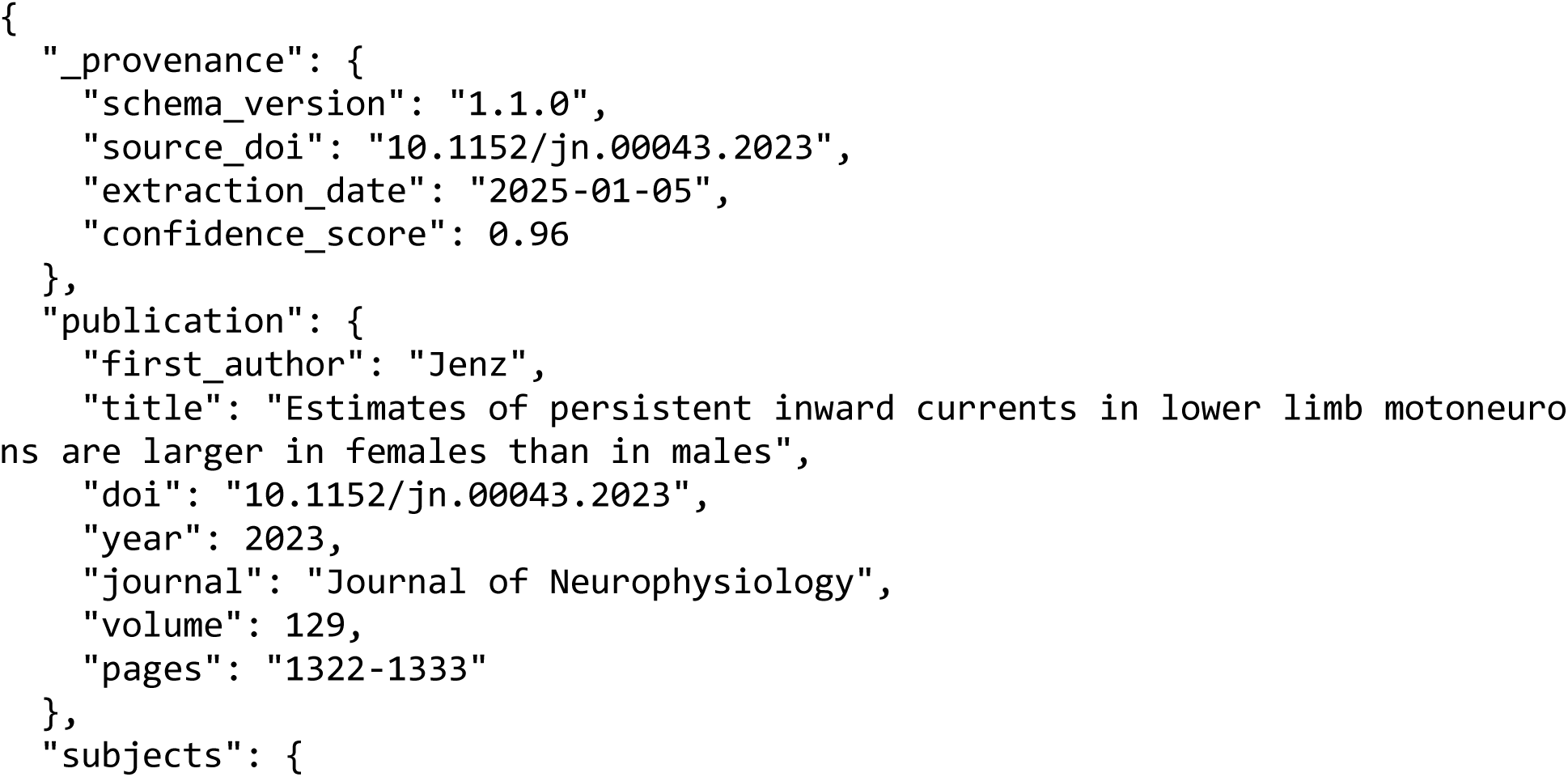
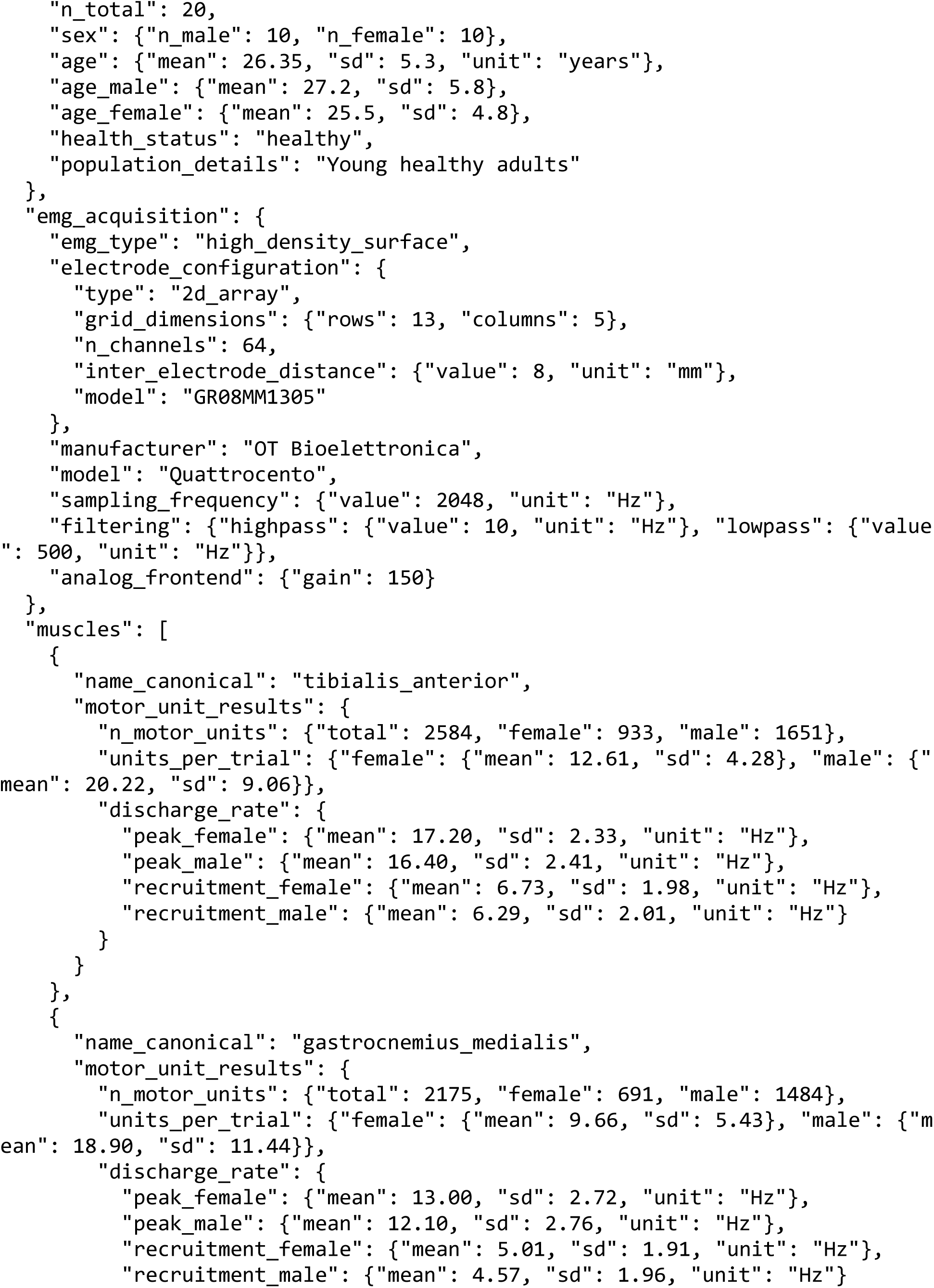
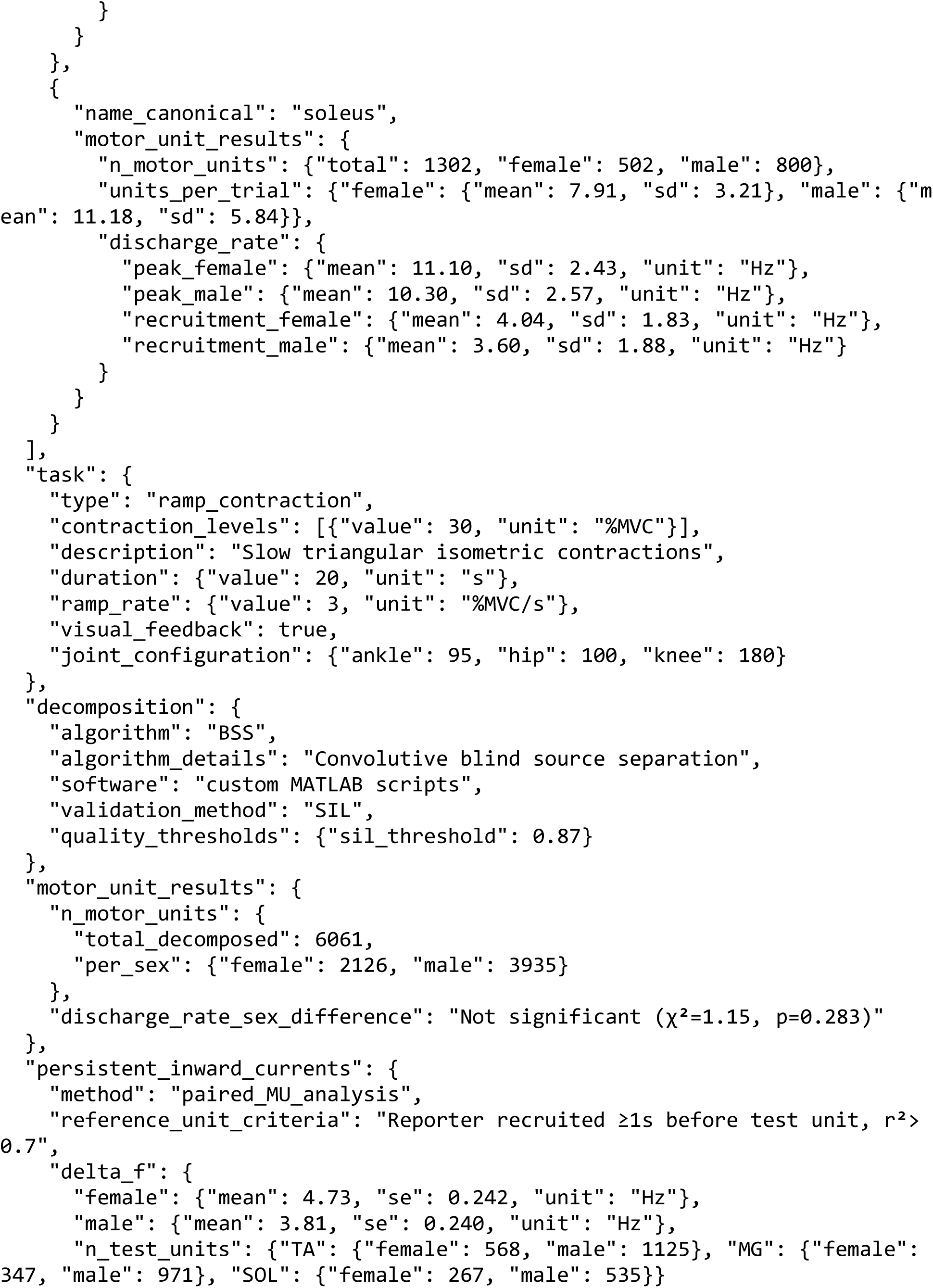
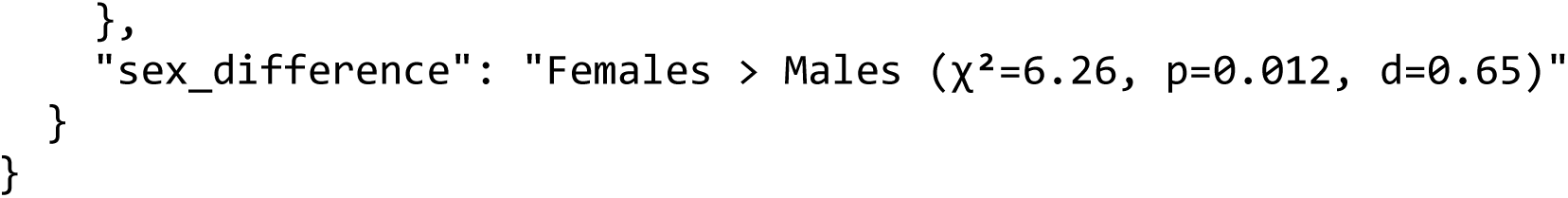

**Primary Schema Sections:**

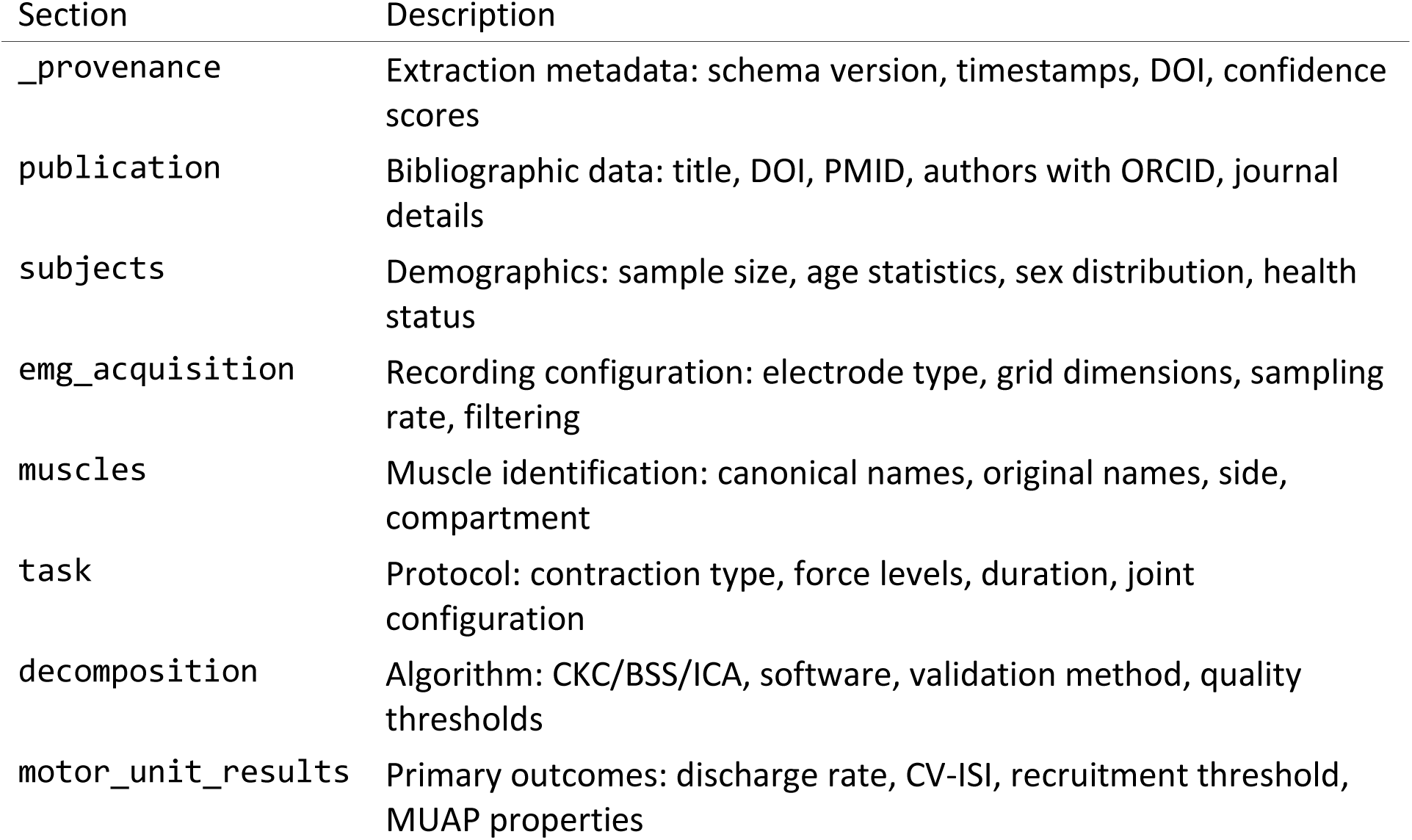

| Section | Description |
| --- | --- |
| _provenance | Extraction metadata: schema version, timestamps, DOI, confidence scores |
| publication | Bibliographic data: title, DOI, PMID, authors with ORCID, journal details |
| subjects | Demographics: sample size, age statistics, sex distribution, health status |
| emg_acquisition | Recording configuration: electrode type, grid dimensions, sampling rate, filtering |
| muscles | Muscle identification: canonical names, original names, side, compartment |
| task | Protocol: contraction type, force levels, duration, joint configuration |
| decomposition | Algorithm: CKC/BSS/ICA, software, validation method, quality thresholds |
| motor_unit_results | Primary outcomes: discharge rate, CV-ISI, recruitment threshold, MUAP properties |

The schema is aligned with BIDS-BEP042 standards for EMG data and includes validation rules for data types, ranges, and required fields.

### S2: Complete EMG Metadata Schema and Controlled Vocabulary

#### S2.1: EMG Metadata Schema Structure

The MUscraper extraction pipeline uses a comprehensive JSON Schema containing **over 200 fields** organized into **17 major sections**, aligned with the BIDS-BEP042 specification for EMG data.

**Extended Schema Sections:**

| Section | Fields | Description |
| --- | --- | --- |
| _provenance | 7 | Extraction metadata: schema version, timestamps, DOI, confidence |
| publication | 12 | Bibliographic data: title, DOI, PMID, authors, journal details |
| subjects | 10 | Demographics: sample size, age, sex distribution, health status |
| emg_acquisition | 25+ | Recording: electrode type, grid dimensions, sampling rate, filtering |
| muscles | 15+ | Muscle identification: canonical names, side, compartment |
| task | 10 | Protocol: contraction type, force levels, duration |
| decomposition | 8 | Algorithm: CKC/BSS/ICA, software, validation (SIL, PNR) |
| motor_unit_results | 40+ | Primary outcomes: discharge rate, CV-ISI, recruitment threshold |
| persistent_inward_currents | 12 | PIC/delta-F estimates from paired motor unit analysis |
| common_synaptic_input | 15 | Coherence, cross-correlation, frequency band analysis |
| neurophysiology_extended | 10 | H-reflex, M-wave, voluntary activation measures |
| twitch_properties | 8 | Spike-triggered averaging: twitch force, contraction time |
| force_control | 12 | MVC, force steadiness (CoV), rate of force development |
| myoelectric_control | 30+ | Prosthetics: classifier type, DoF, accuracy |
| spatial_properties | 8 | Innervation zone location, motor unit territory size |
| fatigue | 10 | Time to task failure, MVC decline, discharge rate changes |
| pathology | 12 | Clinical populations: condition, severity scales |
| intervention | 6 | Training studies: type, duration, frequency, design |
| _extraction_metadata | 8 | Source sections, extraction method, warnings, review flags |

**Data Validation Features:**

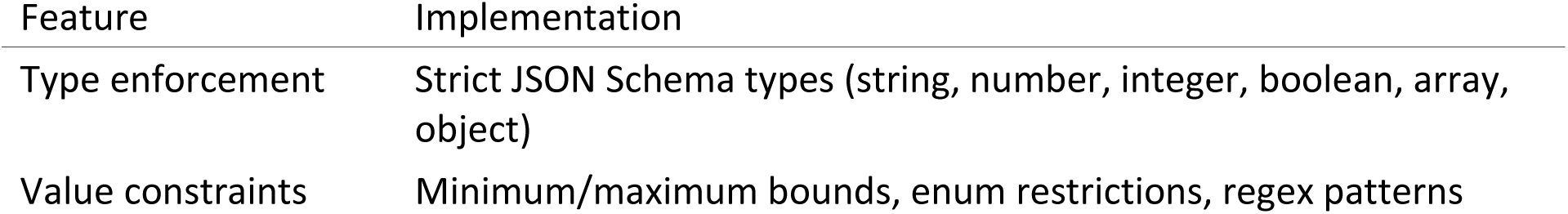
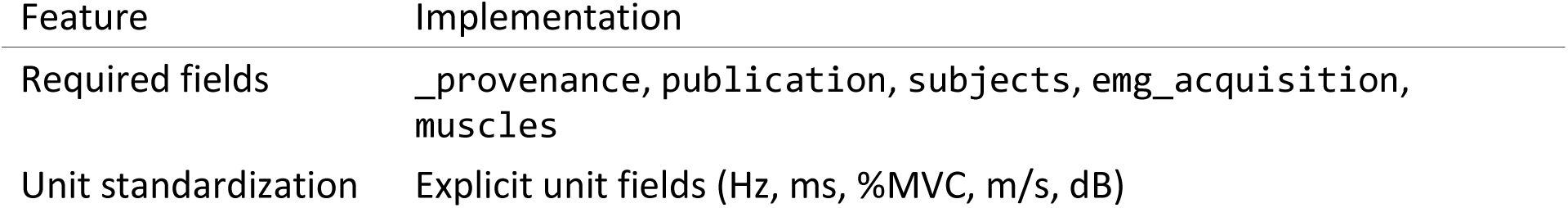

#### S2.2: Controlled Vocabulary

The extraction pipeline uses a controlled vocabulary aligned with BIDS-BEP042 standards to ensure consistent terminology across the corpus.

**Core Motor Unit Terms:**

| Canonical Term | Abbreviations | Units | Description |
| --- | --- | --- | --- |
| `discharge_rate` | DR, FR, MFR | Hz/pps | Motor unit firing rate |
| `instantaneous_discharge_rate` | IDR, IFR | Hz/pps | Instantaneous firing rate |
| `interspike_interval` | ISI, IDI | ms | Interval between discharges |
| `coefficient_of_variation_isi` | CV_ISI | - | Discharge variability |
| `recruitment_threshold` | RT | %MVC | Force at motor unit onset |
| `derecruitment_threshold` | DT | %MVC | Force at motor unit offset |
| `motor_unit_conduction_velocity` | MUCV, MFCV | m/s | Propagation velocity |
| `persistent_inward_current` | PIC, dF | Hz | Estimated motoneuron PIC |
| `common_synaptic_input` | CSI | - | Common drive measure |
| `effective_neural_drive` | END, CST | pps | Cumulative spike train |

**Recording Modality Terms:**

| Canonical Term | Abbreviations | BIDS Field |
| --- | --- | --- |
| `high_density_surface_emg` | HD-sEMG, HDsEMG | EMGChannelType |
| `intramuscular_emg` | iEMG | EMGChannelType |
| `inter_electrode_distance` | IED | EMGElectrodeSpacing |

**Decomposition Quality Metrics:**

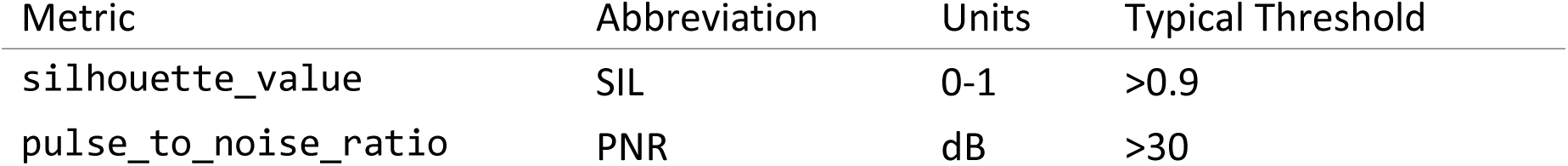

**Muscle Taxonomy (60+ muscles):**

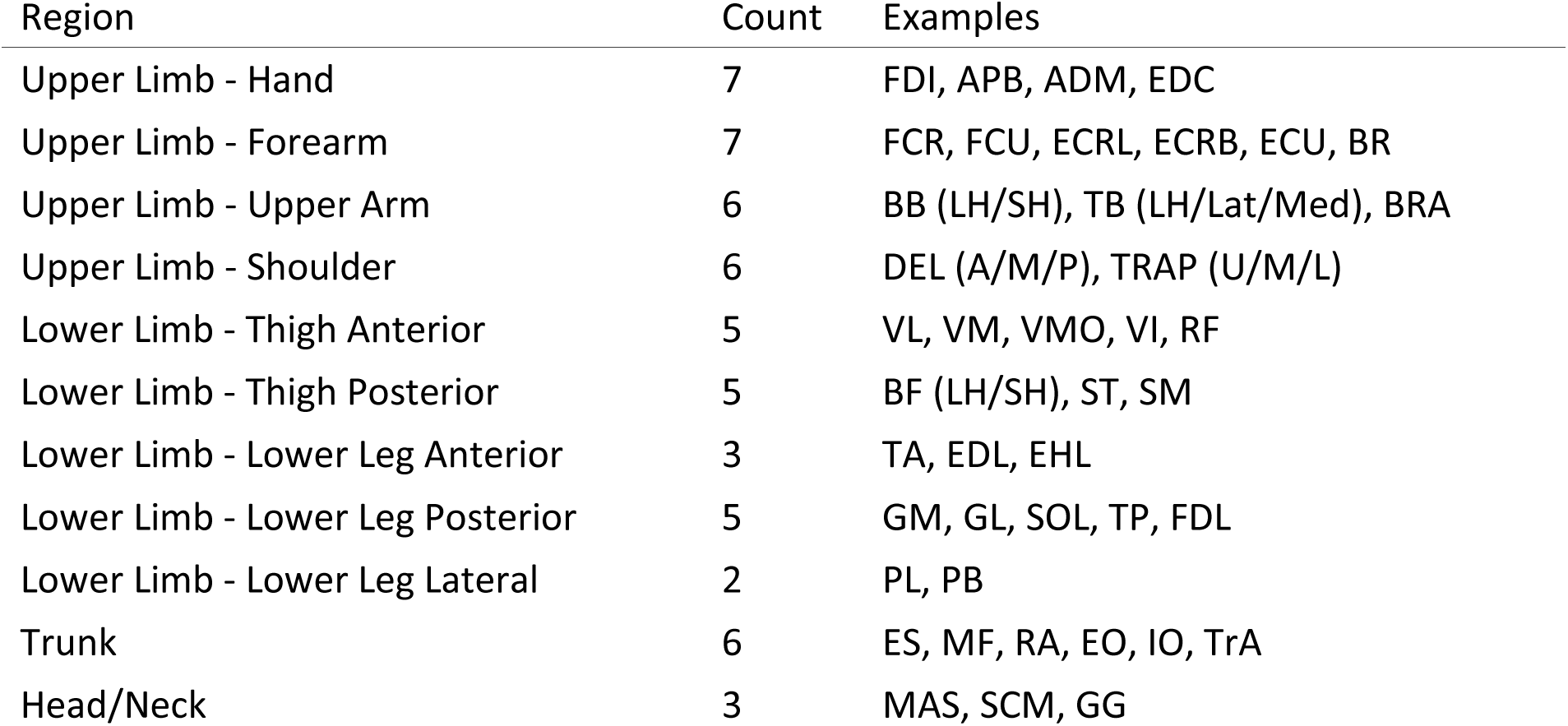

**Muscle Entry Structure Example (Tibialis Anterior):**

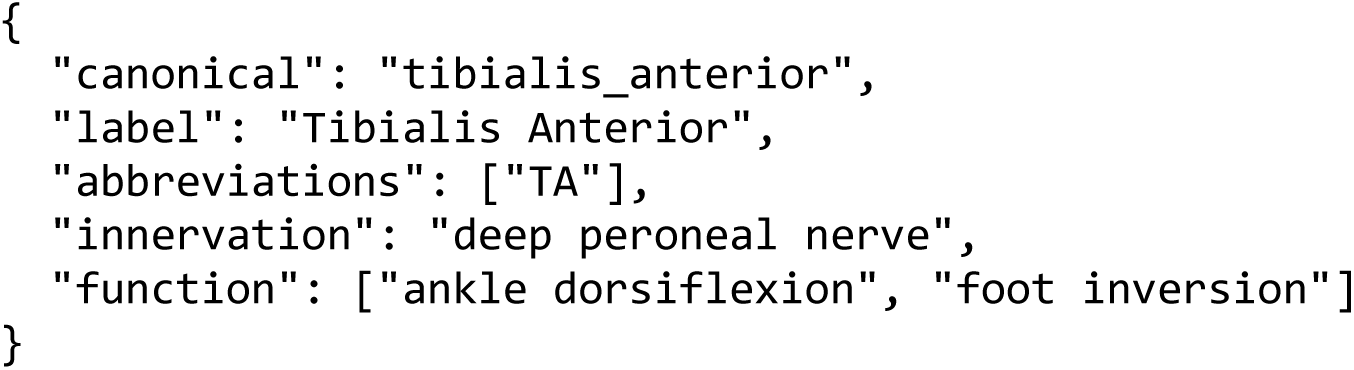

**Myoelectric Control Vocabulary:**

| Category | Terms |
| --- | --- |
| Algorithms | pattern_recognition, proportional_control, regression_control, motor_unit_control |
| Classifiers | LDA, SVM, kNN, MLP, CNN, RNN, LSTM, transformer |
| Features (Time) | MAV, RMS, WL, ZC, SSC, VAR, IEMG |
| Features (Frequency) | MNF, MDF, PKF, MNP, SM |
| Prosthetic Types | transradial, transhumeral, partial_hand, transtibial, transfemoral |

**Equipment Vocabulary:**

| Manufacturer | Products | Type |
| --- | --- | --- |
| OT Bioelettronica | Sessantaquattro, Quattrocento, Muovi, Muovi+ | HD-sEMG |
| Delsys | Trigno, Trigno Avanti, Bagnoli | EMG |
| TMSi | SAGA, Porti | HD-sEMG |

#### S2.3: Schema Usage in MUscraper Pipeline

The schema and vocabulary serve three critical functions:

1. **LLM Prompt Engineering**: The schema structure is provided to the LLM as a template, guiding extraction of structured metadata from unstructured PDF text.
2. **Validation**: Extracted JSON is validated against the schema using JSON Schema Draft 2020-12 validators, ensuring type correctness, required fields, and value constraints.
3. **Harmonization**: The controlled vocabulary enables synonym resolution (e.g., “firing

rate” → “discharge_rate”, “TA” → “tibialis_anterior”), ensuring consistent terminology across the 2,331-publication corpus.

**Validation Statistics:**

- Schema validation pass rate: 94.2% of extracted records
- Common failure modes: missing required fields (3.1%), type mismatches (1.8%), out-of-range values (0.9%)
- Manual review flagged: 12.4% of records for ambiguous or incomplete source text

### S3: MUchatEMG Interface and Representative Outputs

#### S3.1: MUchatEMG Landing Page

**Supplementary Figure S1.**
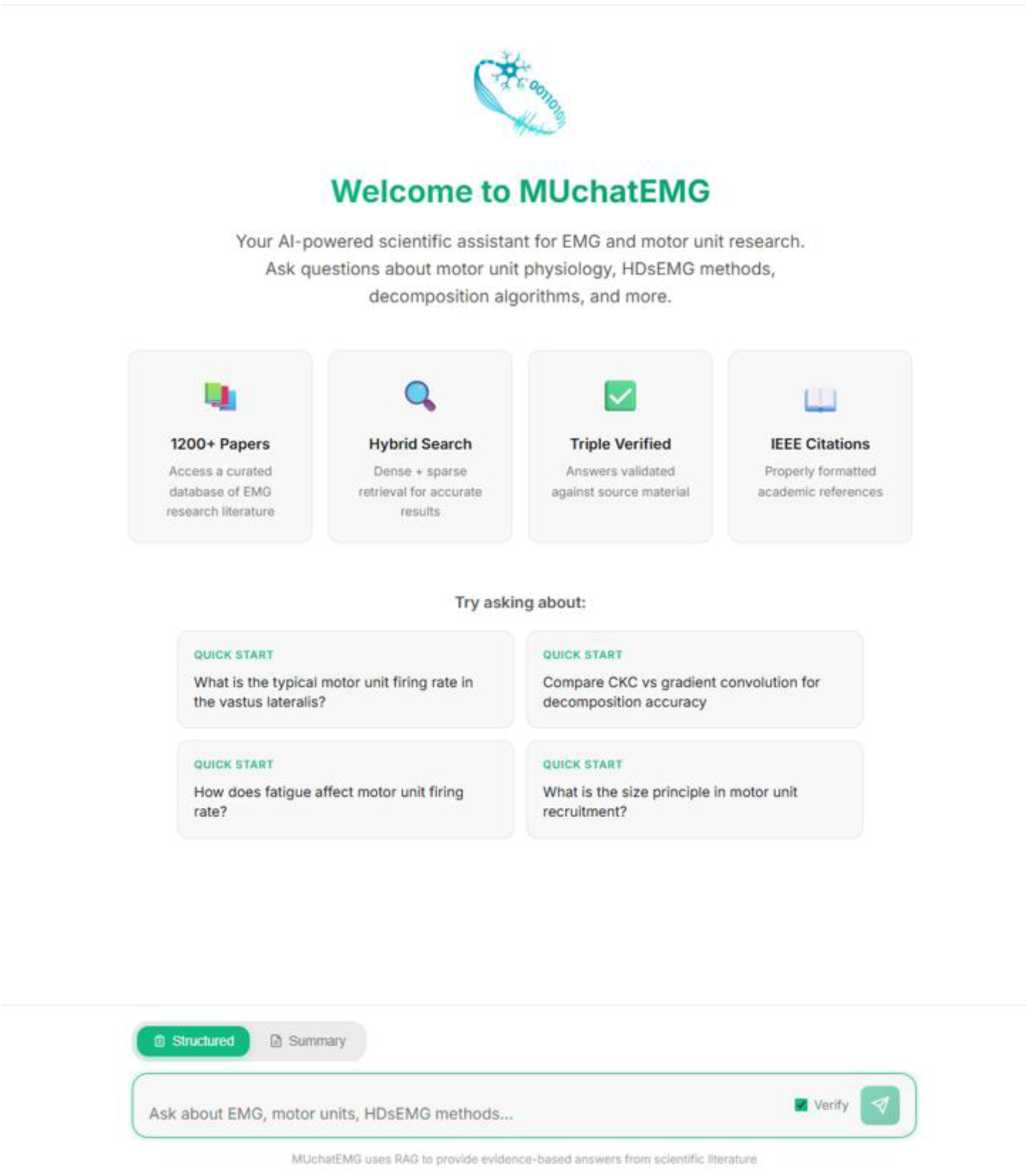
The MUchatEMG web interface. Landing page of the MUchatEMG text summarization system (https://neuro-mechanix.com/). The interface provides access to the curated corpus of over 2,000 peer-reviewed publications on motor-unit and EMG research. Four summary statistics are displayed: the total number of indexed papers, corpus curation and scoring status, topic verification status, and the number of citation-anchored references available. Users can enter natural-language queries in the text field, and MUchatEMG returns structured, citation-grounded responses organized into five tiers of evidence (see Supplementary Tables S3-S4 for examples).

#### S3.2: Representative MUchatEMG Queries and Responses

The following two examples illustrate the structured response format of MUchatEMG. Each response is organized into five tiers of decreasing evidentiary strength, indicated by star ratings:

(1) Grounded Facts [★★★★★]—direct quantitative findings with specific citations; (2) Synthesized Findings [★★★★☆]—cross-study convergences and divergences; (3) Evidence Gaps [★★★⋆⋆]—identified limitations in the current literature; (4) Extended Reasoning [★★⋆⋆⋆]—inferences based on general physiological principles; and (5) Testable Hypotheses [★⋆⋆⋆⋆]—falsifiable predictions with suggested experimental approaches. All citations link to specific peer-reviewed publications in the curated corpus.

**Supplementary Table S3.**
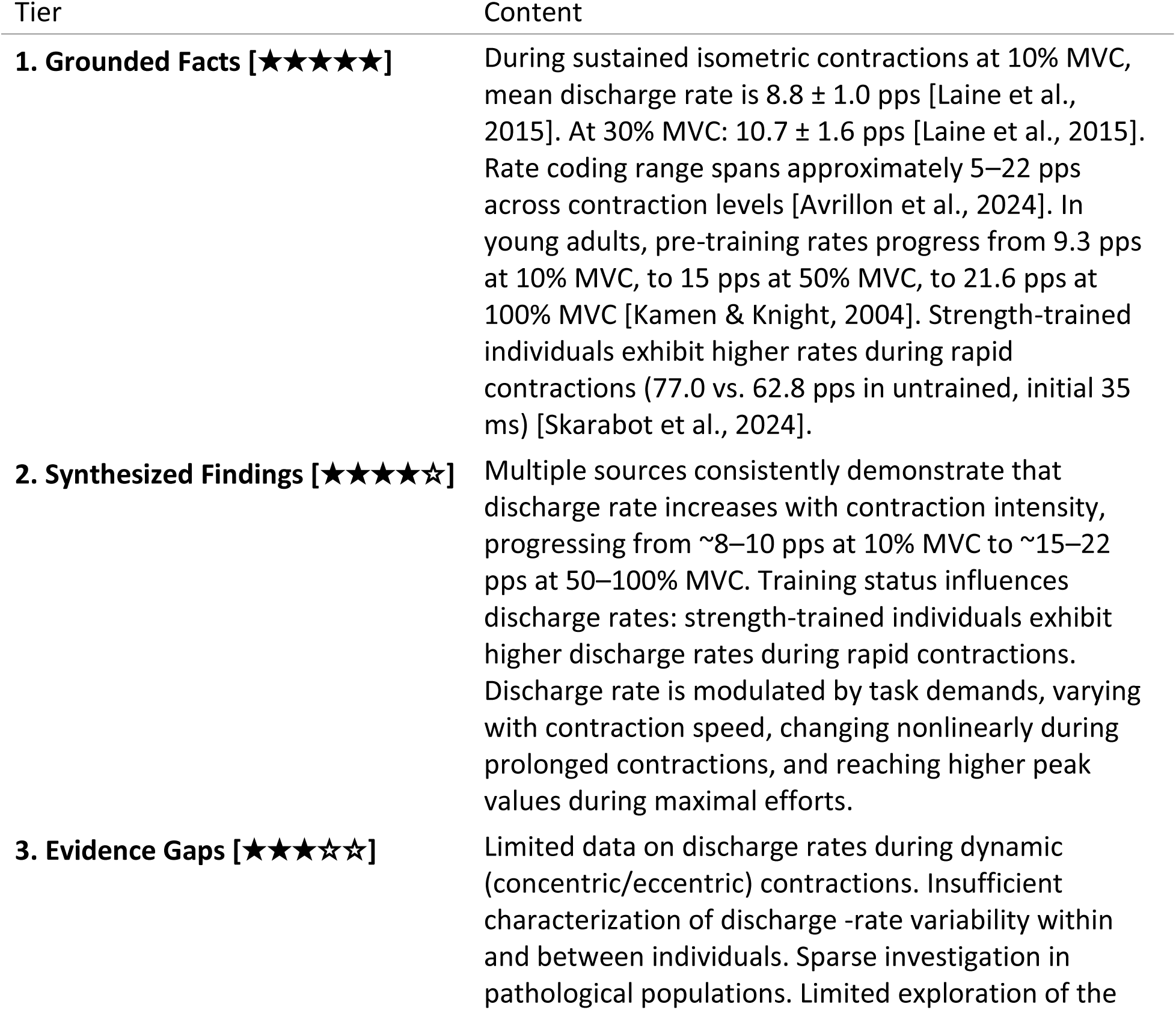
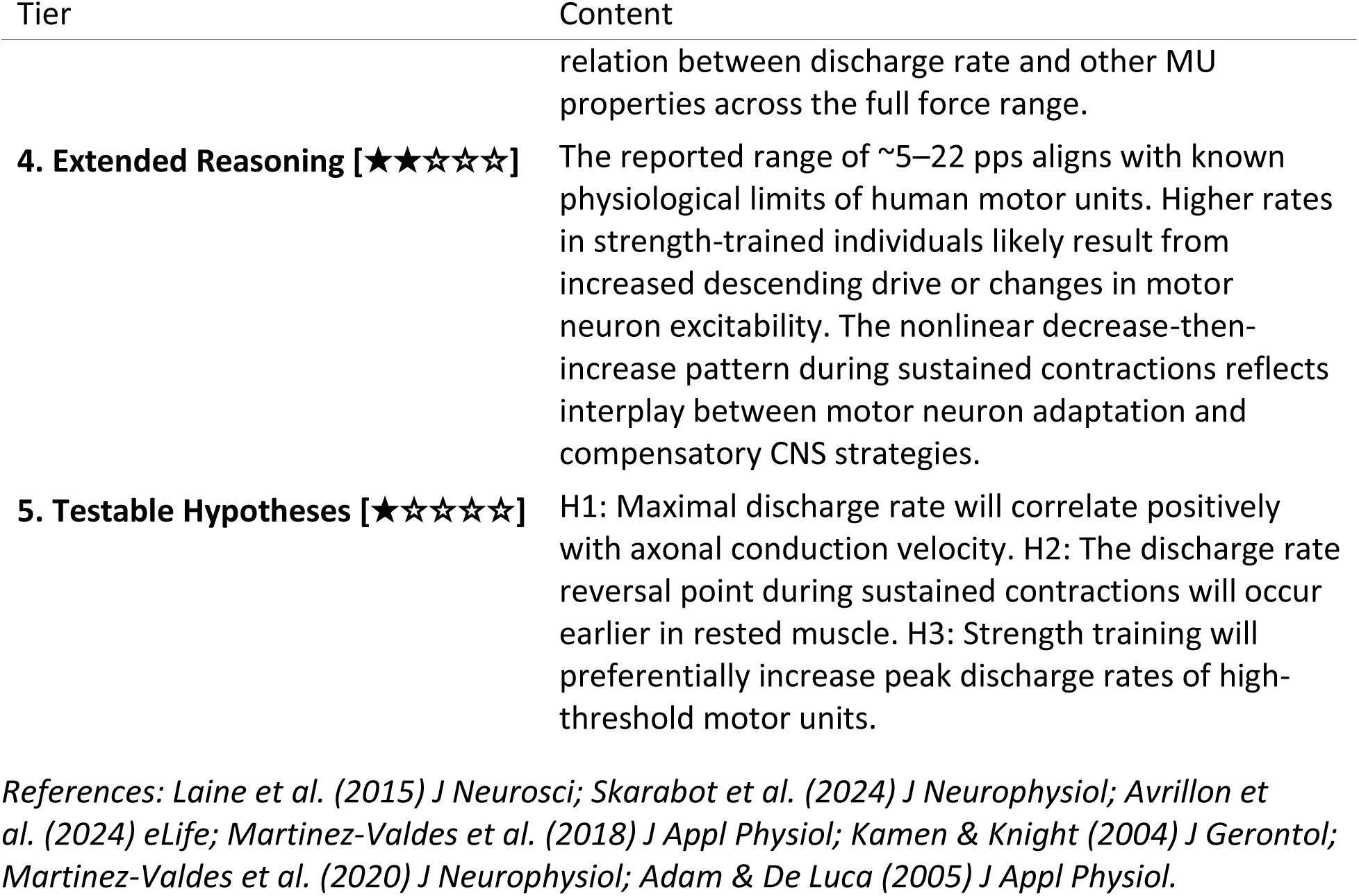
MUchatEMG response to the query: “What is the typical motor-unit discharge rate in the vastus lateralis?”

**Supplementary Table S4.**
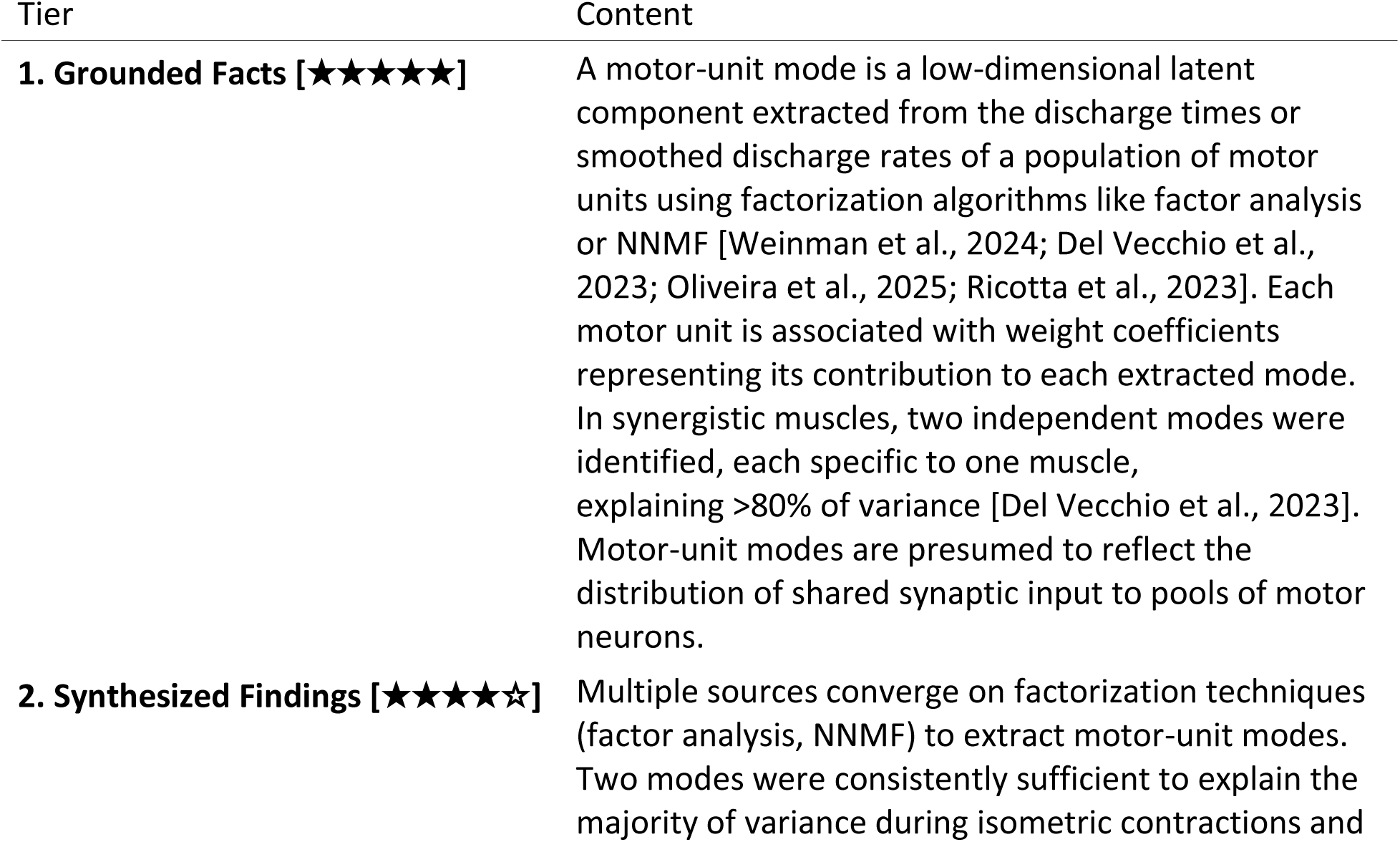
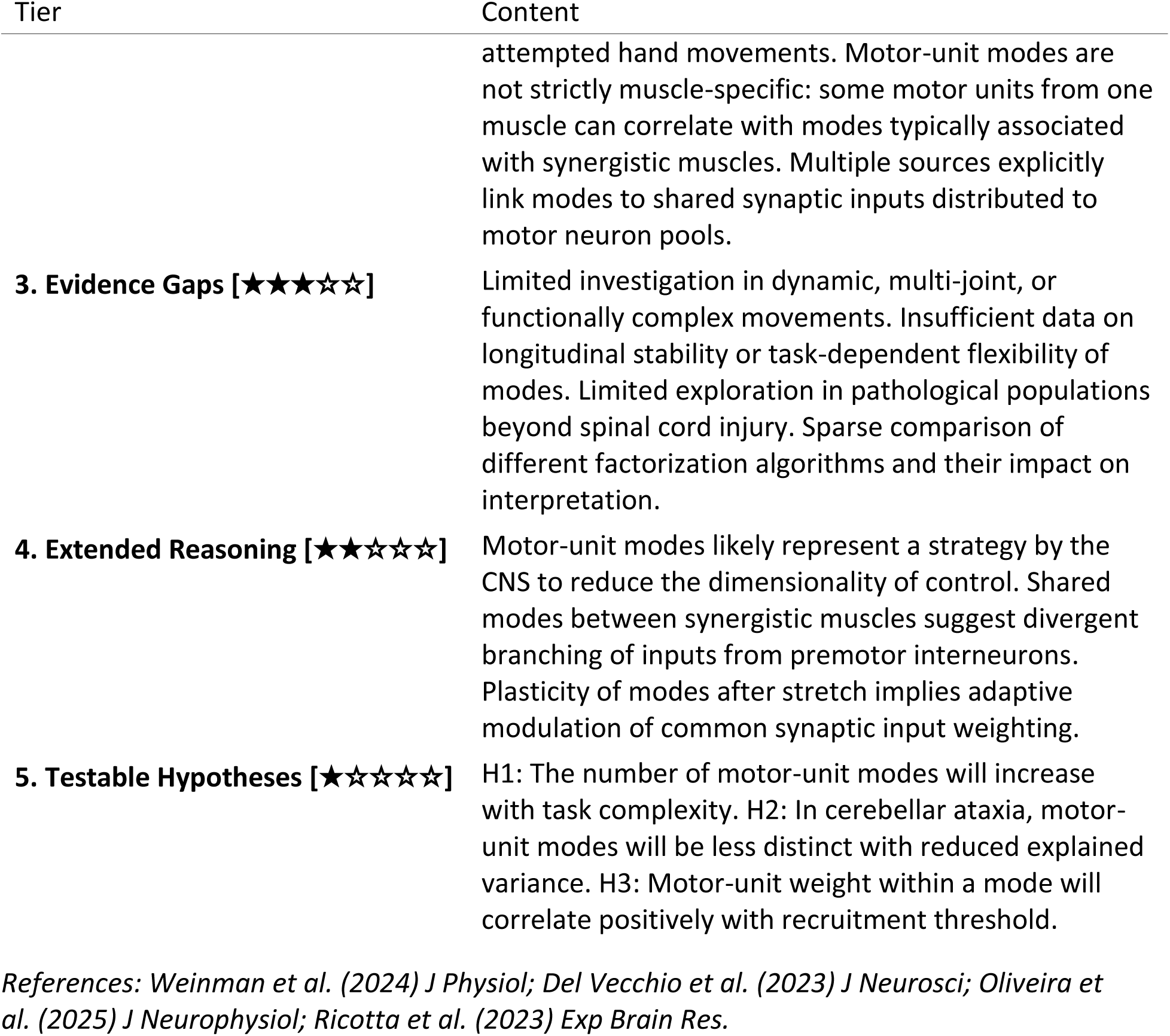
MUchatEMG response to the query: “What are motor-unit modes?”

